# Self-renewal of double negative 3 (DN3) early thymocytes allows for thymus autonomy but compromises the β-selection checkpoint

**DOI:** 10.1101/2020.06.09.142596

**Authors:** Rafael A. Paiva, António G. G. Sousa, Camila V. Ramos, Mariana Ávila, Jingtao Lilue, Tiago Paixão, Vera C. Martins

## Abstract

T lymphocyte differentiation in the thymus relies on high cellular turnover, and cell competition enforces thymocyte replenishment. If deprived of competent progenitors, the thymus can maintain thymopoiesis autonomously for several weeks but this bears a high risk of leukemia. Here we show that double negative 3 early (DN3e) thymocytes can acquire stem cell like properties, which enables them to maintain thymopoiesis. Specifically, DN3e proved to be long-lived, they proliferated and differentiated *in vivo*, were necessary for autonomous thymopoiesis, and included DNA-label-retaining cells. Single cell RNAseq revealed a transcriptional program of thymopoiesis similar in autonomy and the controls. Nevertheless, a new population was identified in thymus autonomy that was enriched for an aberrant Notch target gene signature and bypassed the β−selection checkpoint. In sum, DN3e have the potential to self-renew and differentiate *in vivo* if cell competition is compromised but this enables the accumulation of atypical cells, probably leading to leukemia.

## INTRODUCTION

Normal T lymphocyte development relies on the continuous import of hematopoietic progenitors from the bone marrow. These enter the thymus, commit to the T lymphocyte lineage and go through a series of well-characterized differentiation stages before they finally become T lymphocytes^1, 2^. The time spent at each differentiation stage and the progression between stages is associated with short proliferative phases and prominent cell death^3^. This is in line with the general view that thymocyte developmental stages are generally short and progress through differentiation without self-renewal. Indeed, for a long time, the thymus was considered to lack cells with self-renewal capacity^4^. This was based on classical experiments that showed a transient wave of thymopoiesis from wild type thymi grafted under the kidney capsule of recipients that have an early block in lymphocyte development^5^. However, our work^6^ and that of others^7^ have demonstrated that under specific conditions the thymus can produce T lymphocytes for several weeks independently of bone marrow contribution – thymus autonomy. The implication is that the thymus indeed contains cells that are capable of self-renewal^6, 7^. Specifically, thymus autonomy could be detected 9-10 weeks after thymus transplantation in all recipients deficient for either *Common Cytokine Receptor Gamma Chain* (*γ*_*c*_) or for *IL-7r alpha* (*IL-7rα)*^6, 7^, the two chains that constitute the interleukin 7 receptor (IL-7r)^6, 7^. We further showed that in the steady state thymus autonomy is inhibited by cell competition^8^ occurring at early stages of T lymphocyte differentiation^9^. When cell competition was impaired, thymus autonomy was triggered and later resulted in T cell acute lymphoblastic leukemia (T-ALL) with an incidence of up to 80% and onset at 15 weeks post-transplantation^8, 10^. T-ALL emerging in this experimental setting closely resembled the human pediatric disease in immunophenotype, genetic and genomic alterations, transcriptome and pathological condition^8^. Here we sought to identify the thymocytes that are capable of persisting in the thymus if cell competition is impaired and that can maintain thymopoiesis independently of bone marrow contribution. We show that double negative 3 (DN3) thymocytes, specifically the DN3 early (DN3e), have the potential to self-renew and maintain thymopoiesis autonomously for several weeks. Furthermore, we found that autonomous thymopoiesis was similar to the same process in the steady state and that thymus structure was preserved. Finally, single cell RNAseq (scRNAseq) for gene expression combined with TCR repertoire analyses showed that differentiation indeed relies on a reduced number of precursors. Nevertheless, this process is prone to errors and a new population of thymocytes was seen in autonomy that shows an aberrant gene expression profile, likely to be the earliest stage towards T-ALL.

## RESULTS

### The most immature thymocytes in thymus autonomy are DN3

We sought to identify the most immature thymocytes that can persist in the thymus following competent progenitor deprivation. For that purpose, we performed kinetic studies following thymus transplantations of wild type newborn thymi into *Rag2*^*-/-*^*γ*_*c*_ ^*-/-*^ or into wild type (control) recipients (Fig. 1a). Host derived progenitors of *Rag2*^*-/-*^*γ*_*c*_^*-/-*^ seeded the thymus grafts but failed to outcompete donor thymocytes, which can sustain thymopoiesis autonomously (Fig. 1b). Consistent with former results^9^, from day 7 post-transplantation onwards we never detected the most immature thymocyte populations of donor early T lineage progenitor (ETP) and double negative 2a (DN2a) thymocytes (supplementary Fig. 1a, b). This suggests that these donor thymocytes simply differentiate following transplantation. The subsequent differentiation stages were progressively replaced by host derived thymocytes that were unable to outcompete donor thymocytes from the DN3 stage onwards (Fig. 1c). In the autonomous grafts, thymocytes of donor origin were detected at all stages of differentiation including and following the DN3, for the 9-week duration of the experiment (Fig. 1c, d). This was in contrast with the results in the controls, for which thymocytes of donor origin were completely replaced by host thymocytes within 4 weeks, as expected (Fig. 1c, d). Indeed, despite the reduced absolute numbers in thymus autonomy, there was a clear DN3 population that corresponded to the most immature stage of differentiation, which persisted in the autonomous thymus grafts (Fig. 1e). Altogether, we could detect all stages of differentiation in the autonomous thymi including and following the DN3, suggesting that DN3 are capable of maintaining thymopoiesis under these conditions.

**Figure 1.**
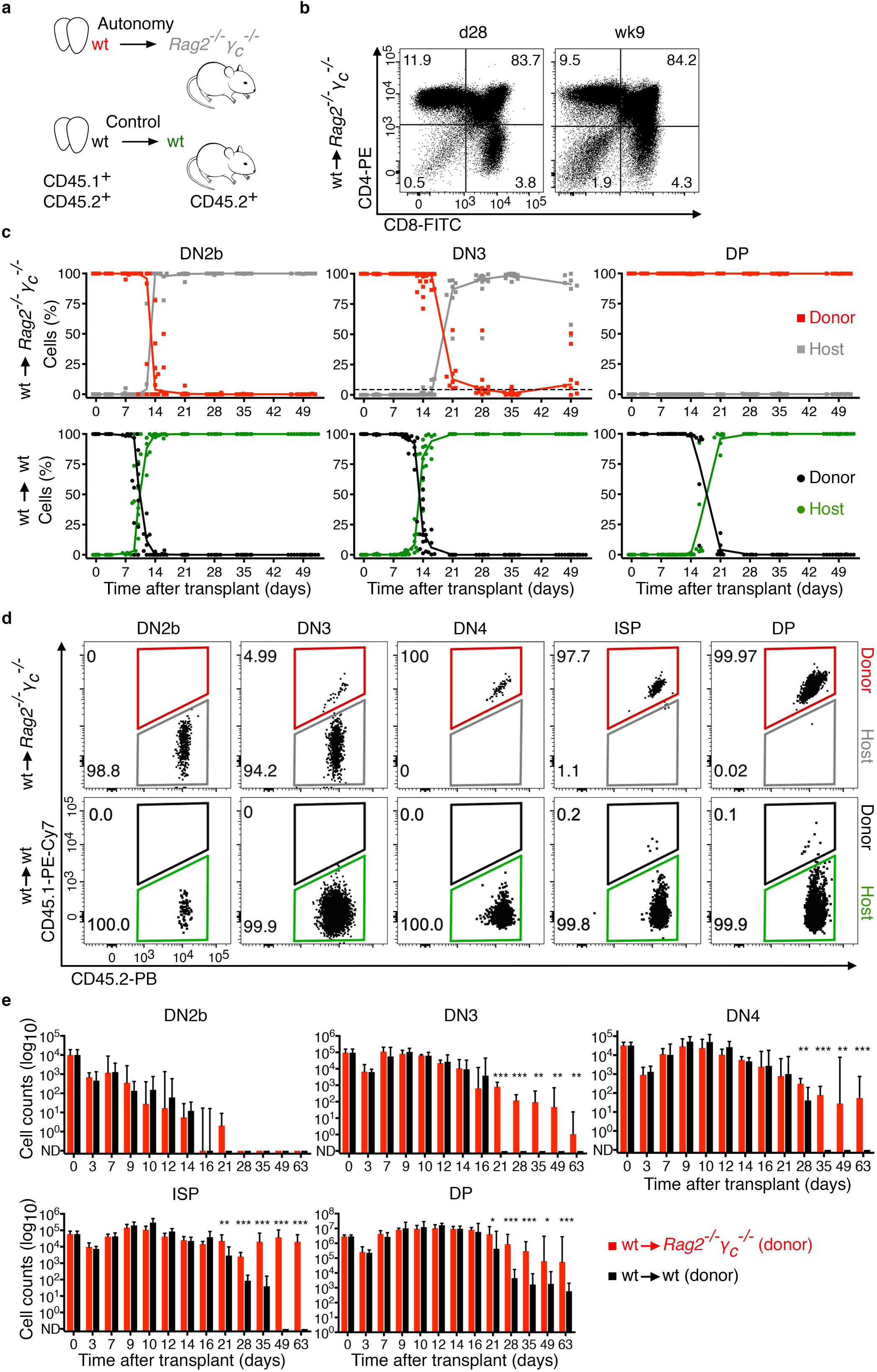
Long-lived DN3 thymocytes revealed by thymus autonomy. **a)** Schematics of the experimental design. Wild type (wt) newborn donor thymi were transplanted into *Rag2*^*-/-*^*γ*_*c*_^*-/-*^ (autonomy) or wild type (control) recipients. **b)** Representative examples of thymus donor thymocytes analyzed 28 days and 9 weeks after transplantation into *Rag2*^*-/-*^*γ*_*c*_^*-/-*^ hosts by flow cytometry. **c)** Percentage of donor or host thymocytes in thymus autonomy (top) or control (bottom) after transplantation for the indicated populations. Lines connect the medians and each pair of dots (for donor and host) represents one graft. Dashed line in the DN3 is the median of the donor DN3 at day 28 (4.8%) **d)** One representative example of thymus autonomy (top) and control (bottom) is shown 28 days after transplant, depicting the donor and host contribution in the indicated populations. **e)** Quantification of the total cellularity of donor thymocytes in the indicated populations over time after transplant. Data was obtained from at least 2 independent experiments with 4 to 12 grafts per timepoint and a minimum of 4 grafts per condition. Data for thymus autonomy (wt into *Rag2*^*-/-*^*γ*_*c*_^*-/-*^) is shown in red (donor) and grey (host) and data for control is depicted in black (wild type donor) and green (wild type host). *p<0.05, **p<0.01, ***p<0.001 Thymocyte populations are DN2b, DN3, CD8+ immature single positive (ISP) and CD4+CD8+ double positive (DP).

### The thymus structure is preserved in thymus autonomy

The thymus is composed of distinct functional regions that are occupied by thymocytes at different stages of differentiation. The most immature stages up to and including the double positive thymocytes locate in the cortex and the most mature CD4 or CD8 single positive thymocytes locate in the thymic medulla. The thymus structure, in particular the 3D network formed by epithelial cells, plays a role in T lymphocyte differentiation. In addition, thymocyte differentiation also impacts on the structure of the thymus. We analyzed wild type thymus grafts upon 4 weeks of autonomy by immunohistology and found that the overall structure and distribution of thymocytes was grossly similar to that of the control thymus grafts (Fig. 2a). The thymus epithelial cells and dendritic cells both had a normal distribution (Fig. 2b). Quantification of the epithelial cells by flow cytometry revealed an alteration in the relative proportions of the different subtypes (Fig. 2c), with slightly more cortical epithelial cells in autonomy, and an increase in the ratio of thymic epithelial cells per thymocyte (Fig. 2d), resulting from the reduction in thymocyte cellularity in the autonomous thymi as compared to the control (not shown). Altogether, only minor alterations could be detected, which were consistent with a normal process of thymopoiesis.

**Figure 2.**
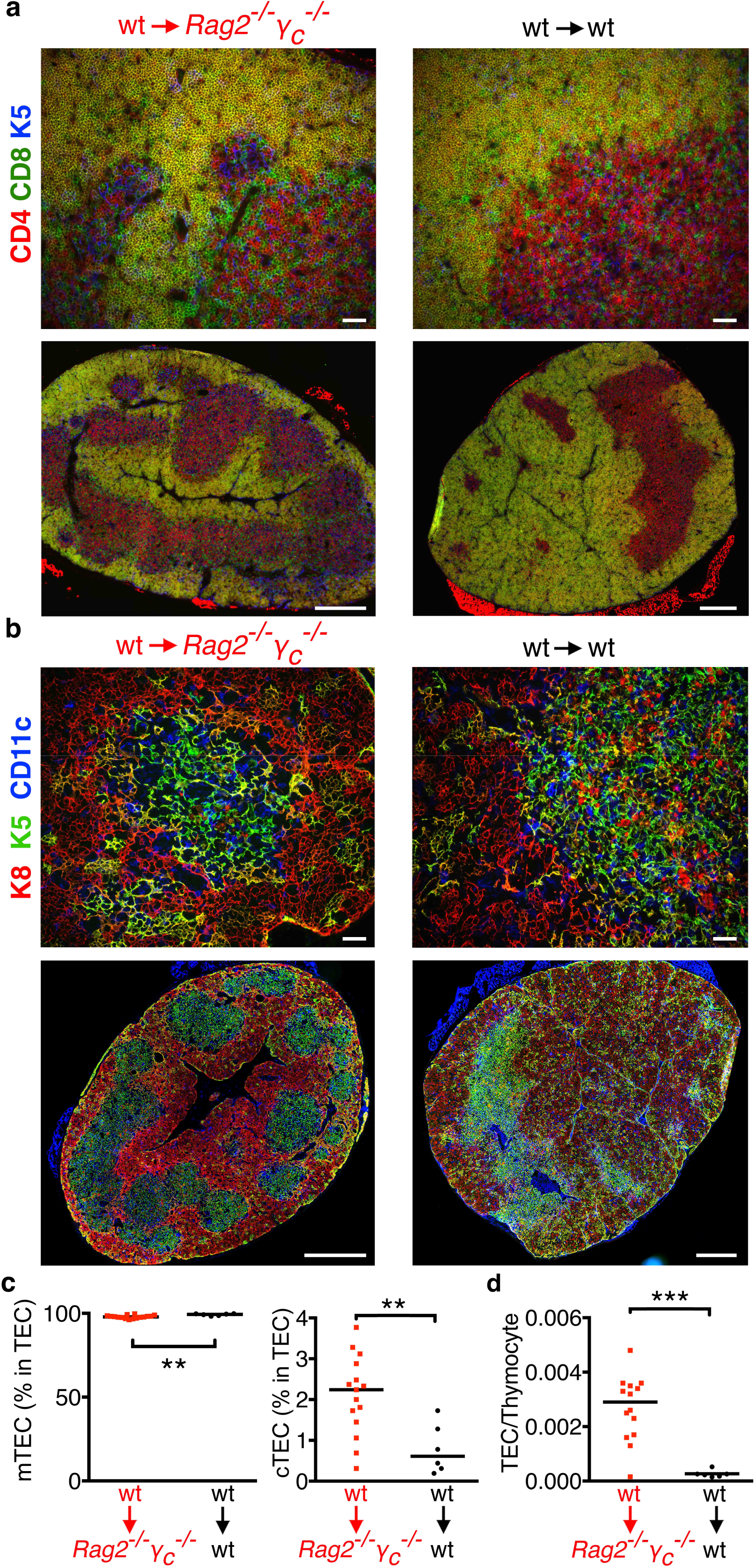
The structure of the thymus is unaltered in thymus autonomy. Wild type (wt) thymi were transplanted into *Rag2*^*-/-*^*γ*_*c*_^*-/-*^ or wild type recipients and analyzed 28 days later by immunohistology (a,b) or flow cytometry (c). **a)** Thymus graft sections were stained for the indicated markers, CD4 (red), CD8 (green) and keratin 5 (K5, in blue), the latter labeling medullary thymic epithelial cells. Yellow shows co-expression of CD4 and CD8. **b)** Sections were stained for the indicated markers, keratin 8 (K8, red), keratin 5 (K5, green) and CD11c (blue) for cortical or medullary thymic epithelial cells and dendritic cells, respectively. Yellow shows co-expression of K5 and K8. Depicted is one representative example of 8 thymus grafts in autonomy and 4 control grafts analyzed. Scale bars = 50 μm (top) and 500 μm in whole graft (bottom). Images were acquired in Leica DMRA2 (top photos) or Nikon High Content Screening (whole-graft), both at a magnification of 20X. **c)** Quantification of the percentage of medullary and cortical thymic epithelial cells (mTEC and cTEC, respectively) from grafts in autonomy or control at day 28 after transplant. **d)** Ratio of cortical thymic epithelial cell per thymocyte in thymus grafts. Data depicted is from 2 independent experiments and each datapoint represents one graft. **p<0.01, ***p<0.001

### DN3 thymocytes maintain thymopoiesis autonomously

Despite the reduction in cellularity, thymus autonomy generated the same thymocyte subsets as in the steady state. Since DN3 were the most immature donor thymocytes persisting in the thymus, we hypothesized that these were the cells responsible for maintaining thymopoiesis. If that were the case, then depletion of DN3 *in vivo* should abrogate thymopoiesis. To test our hypothesis, we injected recipient mice, that had been transplanted 28 days before, with PC61 antibody to deplete DN3 thymocytes (CD25-positive), and analyzed the mice 2 days or 3 weeks after (Fig. 3a). Depletion of DN3 cells was successful (Fig. 3b) and despite some interindividual variability (Fig. 3c), the efficiency of thymus autonomy was reduced (Fig. 3d). Next we sought to determine progenitor-progeny relationships between the different thymocyte populations in thymus autonomy. For that purpose, we performed EdU chase experiments (Fig. 3e) that showed that DN3 and CD8 immature single positive (ISP) thymocytes were the populations with the highest levels of EdU incorporation, indicating that they have a high proliferation rate, while DN4 and double positive thymocytes were less proliferative (Fig. 3f). The label in DN3 was lost within 3 to 4 days, consistent with a fast rate of differentiation into DN4, which acquired the label (Fig. 3f, g). A similar kinetic was observed for ISP, consistent with differentiation into double positive thymocytes, that acquired the EdU label. (Fig. 3f). Focusing on the dynamics of the DN3, the vast majority of the cells were EdU-positive at the beginning of the experiment but they were fast replaced by unlabeled cells, which can be explained by label dilution due to proliferation and/or by replenishment of the compartment by cells that were unlabeled (Fig. 3g). Though the transition transitions from DN3 to DN4, and ISP to double positive, are intuitive to extrapolate from the data, it is more complex to infer directly about the transition from DN4 to ISP because the populations differ significantly in absolute numbers. We developed a mathematical model of EdU incorporation in thymopoiesis for DN3, DN4, ISP and double positive thymocytes and asked about the relative contribution of the processes that could explain the dynamics of EdU label (Fig. 3h). Specifically, we considered that each population could be described with basis on the incorporation of EdU: for each population we had cells that were EdU-positive or EdU-negative. The label could be lost from every population either by differentiation, proliferation (with dilution of EdU), or death. Increase in the percentage of EdU-positive cells could only be explained by differentiation from the previous compartment (Fig. 3h). Model fits to the data were performed using a non-linear least squares procedure. Since the fitting procedure depends on the initial guesses for the values of the parameter, we randomly sampled the parameter space, and the maximum likelihood (post-fit) parameter values were recorded, together with the measurement of their fit quality (Fig. 3i). The parameter estimates for the best-fit models show that even though estimates for the rate of differentiation from DN4 to ISP are broad, they are non-negligible and consistent with normal thymocyte development (supplementary Fig. 2a). This remained true even after restricting the model to consider only values that were consistent with those obtained experimentally, i.e. proliferation rates of DN3 and ISP must be superior to those of DN4 and double positive thymocytes (supplementary Fig. 2b). Lastly, the transition from DN4 to ISP was addressed by comparing the fit-quality of the model for several rates of differentiation and in particular if this developmental transition was removed (k4 = 0). We found that the best fitting considered the transition DN4 to ISP, i.e. k4>0 (Supplementary Fig. 2c). Taken together, the data is consistent with autonomous thymopoiesis being a process that depends on, and progresses from the DN3, thereafter following the same steps as in the steady state.

**Figure 3.**
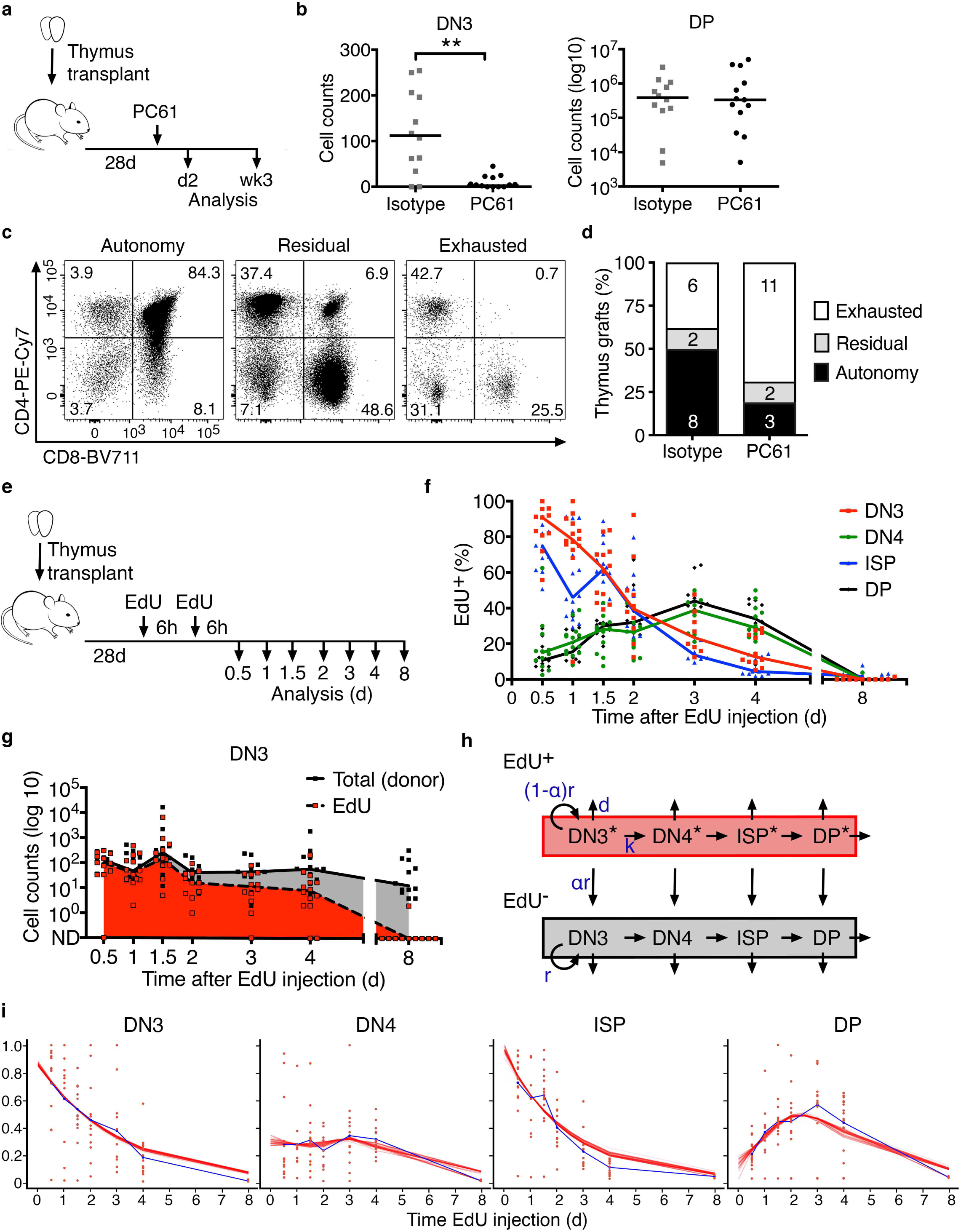
Thymus autonomy depends on DN3 thymocytes. **a)** Schematics of the experimental design. Wild type thymi were transplanted into *Rag2*^*-/-*^*γ*_*c*_^*-/-*^ hosts, which 28 days later were injected i.p. with 250 μg anti-CD25 depleting antibody (PC61) or an isotype control. Analyses were performed 2 days or 3 weeks later. **b)** Thymus grafts were analyzed 2 days after PC61/isotype injection and the indicated thymocyte populations were quantified. **p<0.01 **c)** Thymus grafts were analyzed by flow cytometry 3 weeks after PC61/isotype injection (7-week post-transplant) and the phenotypes obtained for the indicated markers are shown. Thymus grafts with 10% or more CD4+CD8+ double positive (DP) thymocytes were classified as positive for thymus autonomy (left). If DP were under 10%, grafts were classified as residual (middle), or exhausted (right) if no DP population was detectable. **d)** Quantification of the three phenotypes obtained. Numbers in the bars correspond to the number of thymus grafts with that phenotype. **e)** Schematic representation of the experimental design for the EdU pulse-chase assay. Wild type thymi were transplanted into *Rag2*^*-/-*^*γ*_*c*_^*-/-*^ hosts and 28 days later, mice were injected i.p. twice with a 6-hour interval. The first injection was considered time 0 and mice were analyzed at the indicated timepoints. **f)** Quantification of EdU-positive cells in the donor populations, as indicated, over time. **g)** Quantification of donor DN3 thymocytes over time. Total cellularity is shown in black, EdU-positive cell counts are shown in red and the grey shade illustrates the EdU-negative population. Data is from 2 independent experiments. Each symbol represents cells in one graft (red for EdU-positive and black for EdU-negative) and the lines connect the medians. **h)** Schematic representation of the mathematical model considering the proliferation (r) and death (d) rates of each thymocyte population and the differentiation (k) between each developmental stage. **i)** Dynamics of EdU in the indicated populations. Experimental values are the same as in f) depicted as proportions. Dots are values per graft and the blue line connects the mean. Red lines correspond to the estimated dynamics for the top 70 (SSR: [14,14.5]) best-fit parameter combinations. Thymocytes are DN3, DN4, CD8+ immature single positive (ISP) and CD4+CD8+ double positive (DP).

### Persistence of DN3 in thymus autonomy is associated with transcriptional changes and increased proliferation

Since DN3 prolonged their life span and maintained T cell differentiation autonomously for some weeks, we sought to address whether this capacity was associated with changes in whole transcriptome. For that purpose, we sorted total DN3 and double positive thymocytes from autonomous and control thymus grafts and performed bulk RNA sequencing. Principal Component Analysis (PCA) shows that the transcriptional profile of DN3 differed significantly between the two conditions, while differences between double positive thymocytes were substantially less pronounced (Fig. 4a). Specifically, we identified 551 down- and 194 upregulated genes in DN3 in autonomy as compared to their control counterparts (Fig. 4b, Supplementary table 1,2). This was different for double positive thymocytes, for which only 94 and 59 genes which were down- or upregulated in autonomy, respectively (Fig. 4b, Supplementary tables 3,4). Among the upregulated genes in DN3 in autonomy, we found 5 transcription factors (*Tbx21, Arnt2, Ikz3, Dtx1* and *Nfil3*) and genes involved in signaling pathways, namely EGF (*Egfr*), IGF (*Igf2, Igf2bp3*), IFN (*Tbx21, Ctse, Ifi211, Ifi208,Iifi204*), TNF (*Tnfaip2* and *Tnfrsf8*), Wnt (*Sfrp2* and *Wnt5b*) and Notch (*Dtx1* and *Heyl*). Gene ontology analysis of the differentially expressed genes in DN3 (Fig. 4c) detected an enrichment in genes related to cell cycle. Indeed, among the upregulated genes by DN3 in thymus autonomy, 96 were related to proliferation, cell division, metabolism, biogenesis and/or membrane trafficking, consistent with the previous results (Fig. 3). This was consistent with EdU incorporation experiments (Fig. 4d) showing that DN3 proliferate at higher rates in autonomy (Fig. 4e), opposite to all the other stages, which proliferated less than the control counterparts (supplementary Fig. 3a). Similar results were obtained using Ki-67 staining as a proxy for proliferation (supplementary Fig. 3b, c). The increase in proliferation, as well as the many genes involved in metabolism prompted us to test glucose consumption, which was indeed increased in thymus autonomy (Fig. 4f, g). No differences in apoptosis as measured by Annexin V staining were detected between DN3 in autonomy and control thymus grafts (supplementary Fig. 3d). In summary, the transcriptional profile of DN3 changed significantly during autonomy, which reflected the changes in proliferation and associated metabolic requirements.

**Figure 4.**
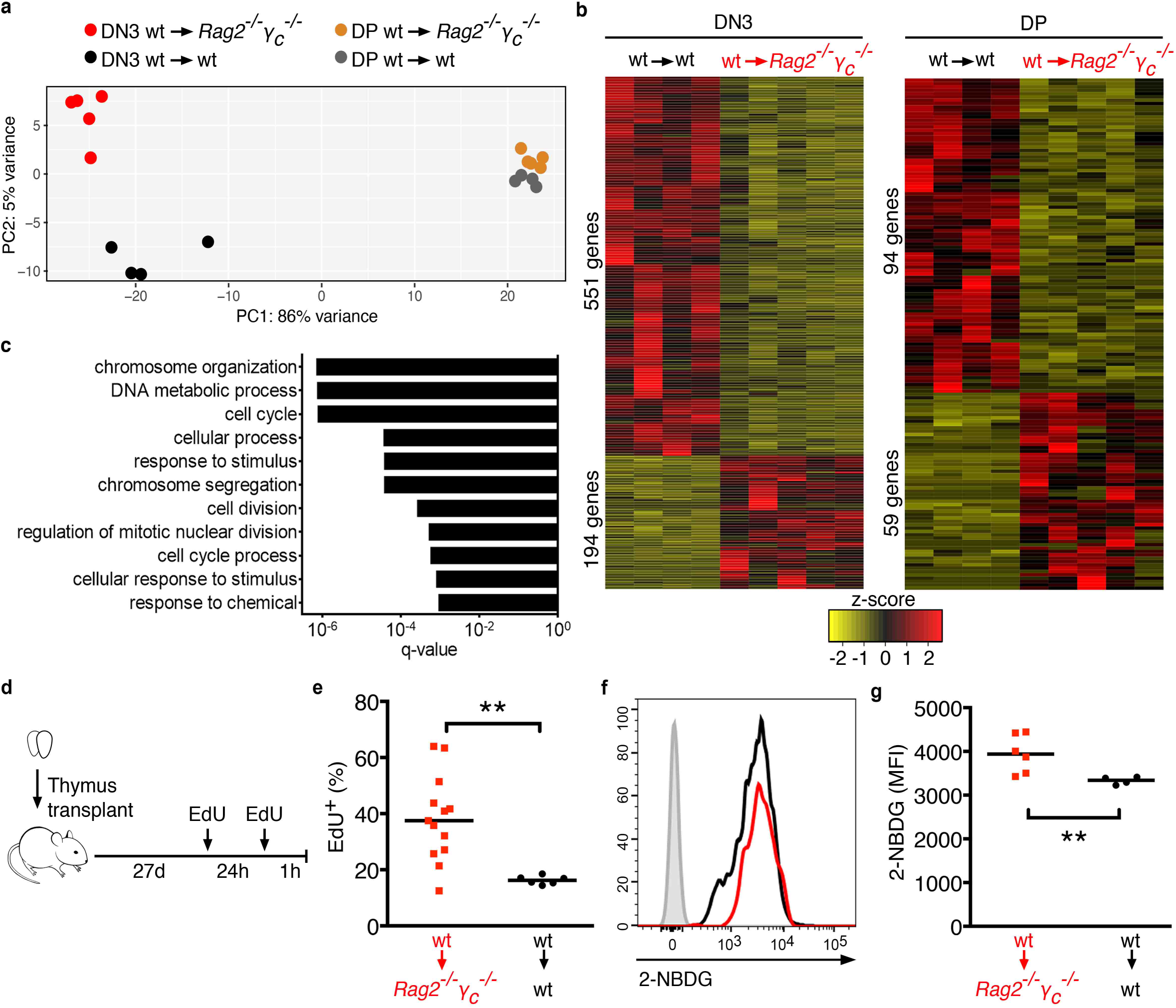
Transcriptomic changes occurring in thymus autonomy. **a)** PCA of bulk RNAseq samples of DN3 and CD4+CD8+ double positive (DP) thymocytes isolated from grafts 28 days after transplant in thymus autonomy (wt into *Rag2*^*-/-*^*γ*_*c*_^*-/-*^) or control (wt into wt). **b)** Heatmaps showing differentially expressed genes between DN3 (left) and DP (right) thymocytes either in autonomy or control with corrected p-value < 0.01 and a fold change of at least 2. **c)** Gene ontology analysis of differentially expressed genes in DN3 in autonomy vs control with corrected p-value <0.05. **d)** Schematic representation of the experimental design for the EdU pulse. Wild type thymi were transplanted into *Rag2*^*-/-*^*γ*_*c*_^*-/-*^ hosts and 27 days later mice were injected i.p. with EdU once, and then again 24 hours later. Mice were analyzed 1 hour after the second injection. **e)** Quantification of EdU-positive thymocytes in donor DN3 in thymus autonomy (red) or control DN3 (black) at day 28 after transplant. **f)** Representative histograms of 2-NBDG incorporation in donor DN3 in thymus autonomy (red) or control DN3 (black), 28 days after transplant. DN3 thymocytes not incubated with 2-NBDG are shown as negative control (grey). **g)** 2-NBDG MFI quantification 28 days after transplant for DN3 in autonomy or control. Each dot represents one graft and the bar corresponds to the median. **p<0.01.

### DN3early self-renew, proliferate more and have more *VDJβ* rearrangements in thymus autonomy

DN3 thymocytes can be subdivided based on the expression of intracellular T cell receptor β (icTCRβ). DN3e do not express icTCRβ while DN3 late (DN3l) do. This transition corresponds to the β−selection checkpoint and depends on the successful rearrangement of the *T cell receptor β (TCRβ)* locus. We found that both subpopulations were present by 28 days of autonomy and their relative proportions were similar to those in the control grafts (Fig. 5a). DN3e were therefore the most immature thymocyte subset detected in thymus autonomy (Fig. 5b). We sought to characterize the DN3 further and assessed the population for the expression of surface markers associated with β−selection, namely the co-stimulatory molecules CD27 and CD28, and the transporters CD71 and CD98^11, 12, 13^. In thymus autonomy, the expression pattern of CD27 and CD28 in DN3 was similar to that of DN3 in the steady state (Supplementary Fig. 4a, c). However, CD71 and CD98 were expressed in DN3e, which does not occur in the steady state (Supplementary Fig. 4b, c), consistent with the changes in proliferation and metabolism of total DN3 (Fig. 4, Supplementary Fig. 2). We evaluated proliferation by EdU incorporation (Fig. 5c) and surprisingly found that DN3e in autonomy proliferated at significantly higher levels than their counterparts in the steady state (Fig. 5d). DN3e thymocytes in the steady-state consist mostly of resting cells undergoing *TCRβ* rearrangement. Therefore, we reasoned that the long persistence and proliferation rate of the DN3e in autonomy was likely to impact on the rearrangement of the *TCRβ* locus. We sorted DN3 at single cell level, aware that the population included approximately 25% DN3l in both conditions. We tested the genomic DNA by nested PCR followed by Sanger sequencing and found that, opposite to the control, DN3 in the autonomous thymi have a higher proportion of *VβDβJβ* and less *DβJβ* rearrangements (Fig. 5f). Intriguingly, while the majority of *VβDβJβ* rearrangements in control DN3 were productive, the opposite was true in autonomy (Fig. 5f). Altogether, these data suggest that DN3e maintaining thymopoiesis autonomously have gone through more cycles of cell division and this is imprinted on the status of their *TCRβ* locus. Finally, we performed BrdU label retention experiments (Fig. 5g) that show that the DN3e included cells that retained label for 14 days after removal of BrdU from the drinking water (Fig. 5h, i). These experiments formally demonstrate that the DN3e in autonomy contain cells with stem cell-like properties, that self-renew to preserve the population while also differentiating to generate their progeny.

**Figure 5.**
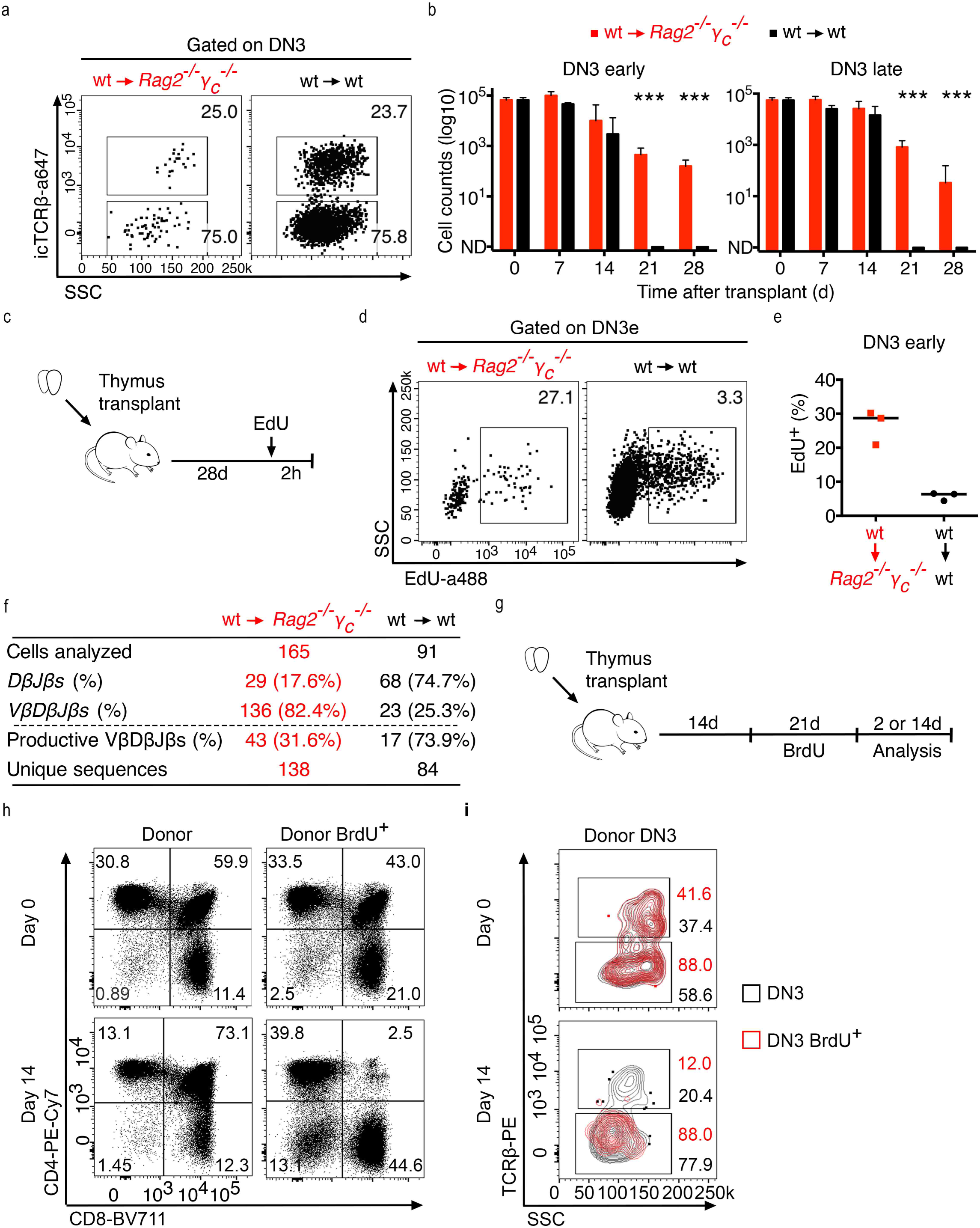
DN3e self-renew, proliferate and have more VDJ rearrangements in autonomy. **a)** Thymus transplants were performed as in Fig.1a and analyzed 28 days later. Cells were gated on donor (left) or total (right) DN3. **b)** Numbers of donor DN3e or DN3l at the indicated timepoints after transplant. Data from at least 2 independent experiments (4-10 grafts per timepoint). ***p<0.001. **c)** Schematics of the experimental design for the EdU pulse. 28 days after thymus transplant, mice were injected i.p. with EdU and analyzed 2 hours later. **d)** Thymus grafts were pooled for analysis (6-8 in autonomy and 2 controls), and depicted are DN3e from one out of 3 independent experiments. **e)** Quantification of the percentage of EdU-positive DN3e in autonomy or control. Data from 3 independent experiments. **f)** Summary of *DβJβ* and *VβDβJβ* rearrangements in DN3, 28 days after transplant. **g)** Schematics of the experimental design. Mice were kept with BrdU in the drinking water (0.8 mg/ml) for 3 weeks starting at day 14 after transplant, and analyzed 2 and 14 days thereafter by flow cytometry. **h)** Depicted are donor thymocytes (left), further gated on BrdU-positive (right) 0 or 14 days after BrdU withdrawal. **i)** Contour plots show donor DN3 thymocytes at day 0 (top) and day 14 (bottom). Black shows DN3 and red are overlaid BrdU-positive DN3. At day 0, 8 individual grafts were analyzed separately, and at day 14, a total of 10 grafts were pooled for analysis.

### Autonomous thymopoiesis assessed by single cell RNA sequencing

Thymopoiesis in autonomy resembled the same process in the steady state, despite progressing from thymocytes that self-renew. We sought to address whether this could be corroborated at single cell level and/or if we could retrieve additional information. For that purpose, we performed single cell RNA sequencing (scRNAseq) and compared thymocytes from 4 thymus grafts in autonomy and one control, which were enriched for the less represented cell populations. Specifically, DN3 to DN4, ISP and double positive thymocytes were sorted in numbers that were similar between the two samples (autonomy and control). A total of 7488 thymocytes encompassing the autonomous and the control thymus grafts could be analyzed (Fig. 6a). These consisted of 11 clusters, which we sought to identify with basis on their gene expression as compared with the available data in ImmGen^14^ and has been further detailed for thymocytes^15^. We found the correspondence for most clusters to known thymocyte populations (Fig. 6b). Specifically, we identified the clusters for thymocytes at the DN3, DN4-ISP, and the different sequential stages within the CD4+CD8+ double positive compartment: blast, small resting, positively selected (CD69-positive) (Fig. 6b). We found two clusters that we could not match to known subsets (clusters 5 and 6) that are close to double positive thymocytes but have a gene expression profile that is suggestive of an activated state. Since the purpose of sorting was to enrich for the small populations, we also purified two additional unexpected populations: mature CD8 T lymphocytes, and myeloid cells (Fig. 6b). The thymocyte distribution was similar between the two conditions (Fig. 6c), and this could be verified by the expression of the top conserved marker genes that defined the clusters (Fig. 6d). Likewise, the expression of selected genes that are associated with thymopoiesis followed the expected pattern from normal T lymphocyte differentiation (Fig. 6 e). However, one additional cluster was detected specifically in thymus autonomy: cluster 4 (Fig. 6b, c). While cells in this cluster resemble double positive thymocytes, their gene expression profile differs sufficiently to define a separate cluster. Altogether, the scRNAseq data was consistent with a differentiation process that was mostly conserved in thymus autonomy, with the expected subpopulations mostly unaltered, but revealed one unexpected cell population, which points at the existence of a parallel program in thymus autonomy.

**Figure 6.**
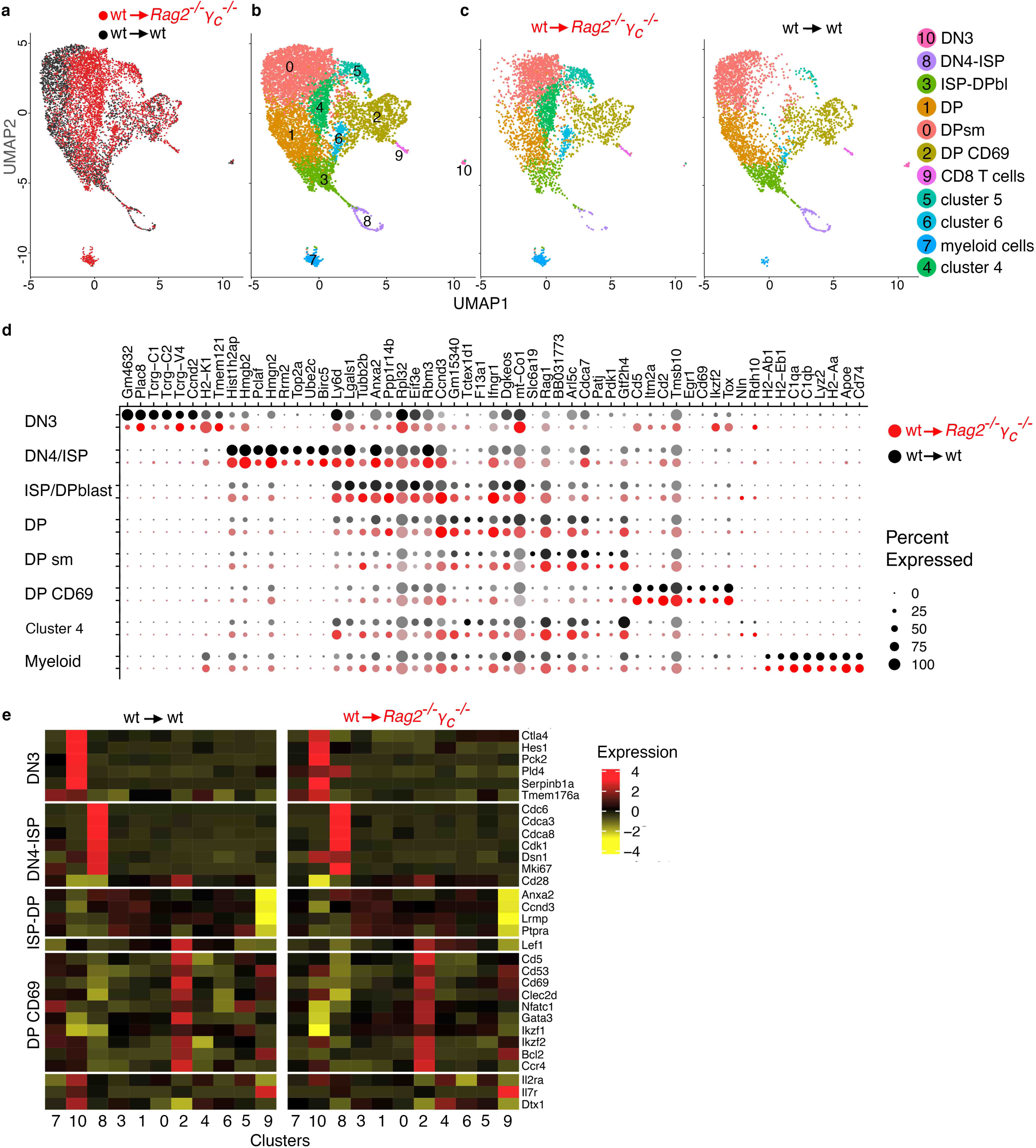
Thymopoiesis in thymus autonomy versus the steady state. DN3, DN4, CD8+ immature single positive (ISP) and CD4+CD8+ double positive (DP) thymocytes were sorted from wild type thymus grafts into *Rag2*^*-/-*^*γ*_*c*_^*-/-*^ (autonomy), or wild type (control) recipients to enrich for the less-represented populations. Cells were analyzed by single cell RNA sequencing (scRNAseq) using the 10X Genomics platform. **a)** UMAP for thymocytes in autonomy (red) and control counterparts (black). **b)** UMAP depicting the different clusters identified and their correspondence to known thymocyte populations (far right). DPbl and DPsm correspond to blast and small resting DP thymocytes, respectively. **c)** Shows the different cluster composition in thymus autonomy (left) and in the control thymocytes (right). **d)** Depicted are up to ten marker genes per cluster, showing the expression in thymus autonomy (red circles) versus control (grey to black circles), where the diameter of the circle represents the percentage of expressing cells that particular gene and the color intensity represents the level of expression. **e)** Heatmap of the mean value of gene expression in the depicted clusters for selected genes.

### Common ancestry in thymus autonomy revealed by thymocytes sharing common *TCRβ*

The scRNAseq experiment was coupled with TCR repertoire analysis, which enabled the identification of cells expressing only *TCRβ* (Fig. 7a) and cells expressing the chains for the complete *TCRαβ* (Fig. 7b). Of note, cluster 2 corresponding to positively selected thymocytes, is highly enriched for cells expressing both *TCRα* and *TCRβ*, and lacking in cells expressing exclusively the *TCRβ* chain (Fig.7a, b). Also surprising was the detection of *TCRα* and *TCRβ* in cluster 7, corresponding to myeloid cells (Fig.7a, b). This most likely results from the mRNA from phagocyted thymocytes in macrophages. When assessing the *Vβ* and *Jβ* usage in rearrangements, it became clear that there was a skewing of the repertoire in thymus autonomy, with preferential usage of distal *Vβ* genes and the *Jβ2* cluster (Fig. 7c). This is consistent with locus erosion possibly due to several rounds of rearrangement. This was not the case in the control thymocytes, where turnover promotes replenishment of thymocytes and *Vβ* and *Jβ* usage is thereby well-distributed (Fig. 7c). When assessing the *TCRα* locus, no skewing was evident in neither of the samples (Fig. 7d). The 50 most prevalent CDR3 sequences in autonomy expressed only *TCRβ* chains and were found to be represented across the majority of the clusters (Fig. 7e). For each of these 50 CDR3 sequences, we could find several cells expressing one common *TCRβ* chain, and then a substantial number of other cells that in addition expressed a unique *TCRα* chain (Fig. 7f). These data are consistent with a limited number of precursors bearing a rearranged *TCRβ* that self-renew and differentiate to maintain thymus autonomy.

**Figure 7.**
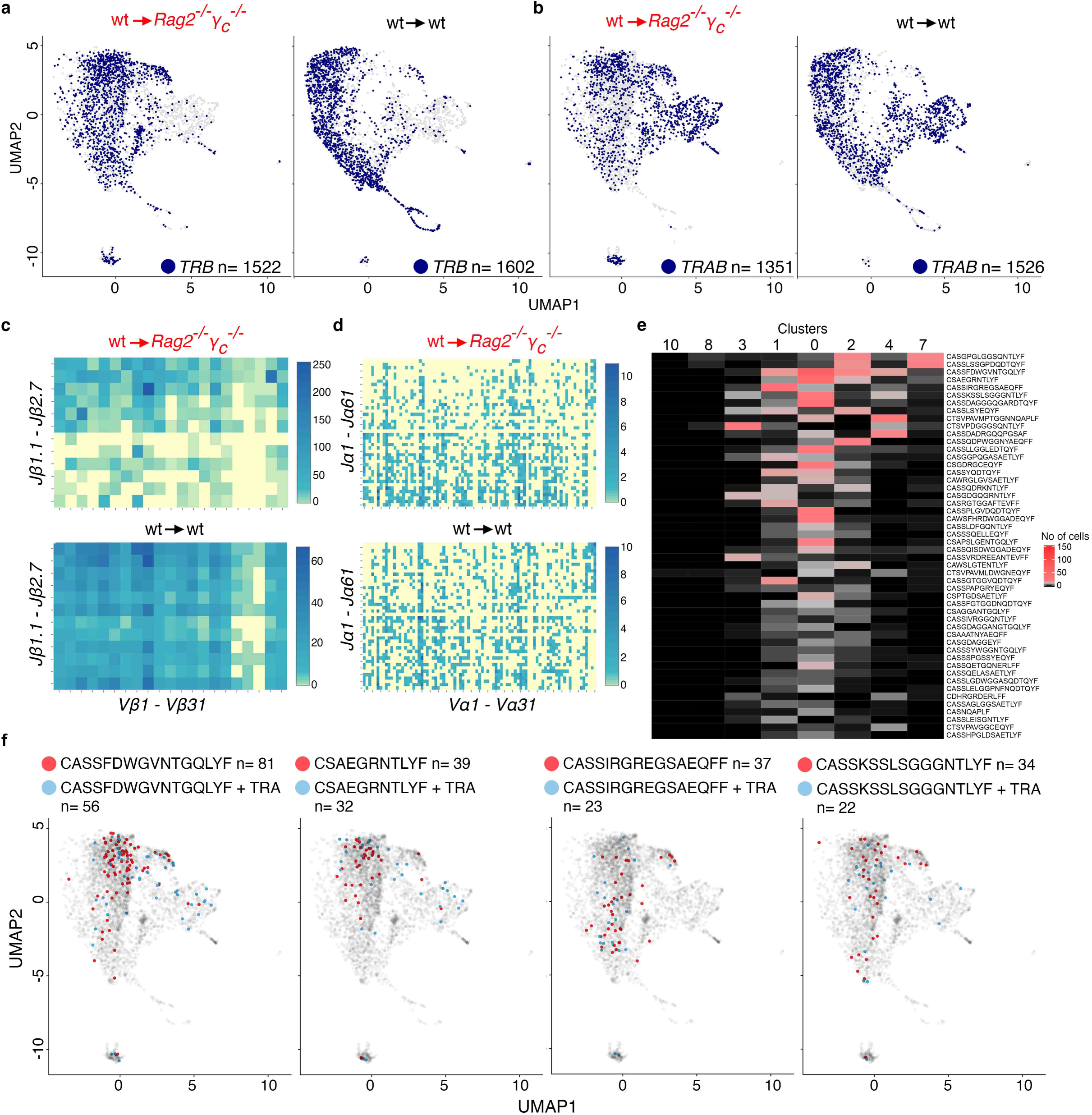
TCR repertoire at the single cell level in thymus autonomy. **a)** UMAPs showing thymocytes with productive *TCRβ* in autonomy (left) or control thymocytes (right). **b)** UMAPs displaying cells with productive *TCRαβ* in autonomy (left) or control thymocytes (right). **c)** Heatmaps representing the number of cells with each possible combination of *Jβ* to *Vβ* in autonomy (top) or control (bottom). Scale is to the right. **d)** Heatmaps showing the number of cells with each possible recombination of *Jα* to *Vα* in autonomy (top) or control (bottom). **e)** Heatmap showing the distribution across clusters for the 50 most prevalent CDR3s *TCRβ* in thymus autonomy. **f)** UMAPs highlighting four examples of prevalent *TCRβ* (red) that is common to several thymocytes. Red correspond to cells only expressing the indicated CDR3 for *TCRβ*. Blue depict cells sharing the same *TCRβ* but each expressing a unique *TCRα.*

### Emergence of a new population in thymus autonomy

Our data support that self-renewing DN3e maintain thymopoiesis that follows a transcriptional program similar to that in the steady state. However, scRNAseq revealed one new population (cluster 4) that would have been undetected by other techniques, and is absent in the steady state (Fig. 6b, c). Although these cells resemble double positive thymocytes, their transcriptional profile segregates them from all the other clusters that are shared with the control sample. Interestingly, thymocytes in autonomy, with a strong prevalence in the cells of cluster 4 express *Notch1* and have an aberrant Notch target gene expression, most notably by expressing *Deltex1* that is barely detectable in control thymocytes in the steady state (Fig. 8a). Furthermore, the same cluster was enriched for cells with failed, non-productive *TCRβ* rearrangements (Fig. 8b). This indicates that those cells overcame the β−selection checkpoint even though they lacked a functional TCRβ. The presence of double positive thymocytes lacking functional TCRβ could be detected by flow cytometry in a substantial number of autonomous thymus grafts (Fig. 8c, d), confirming that such cells indeed emerge in these conditions. This suggests that in parallel with the normal differentiation, there is an additional program that operates in distinct, individual cells potentially creating the conditions for the malignant transformation that later causes leukemia^8, 10^. Altogether, we could identify an aberrant gene expression program running in parallel with normal thymopoiesis that was characterized by thymocytes escaping the β-selection checkpoint and expressing an unusual pattern of Notch target genes, potentially the earliest cells on the path to T-ALL.

**Figure 8.**
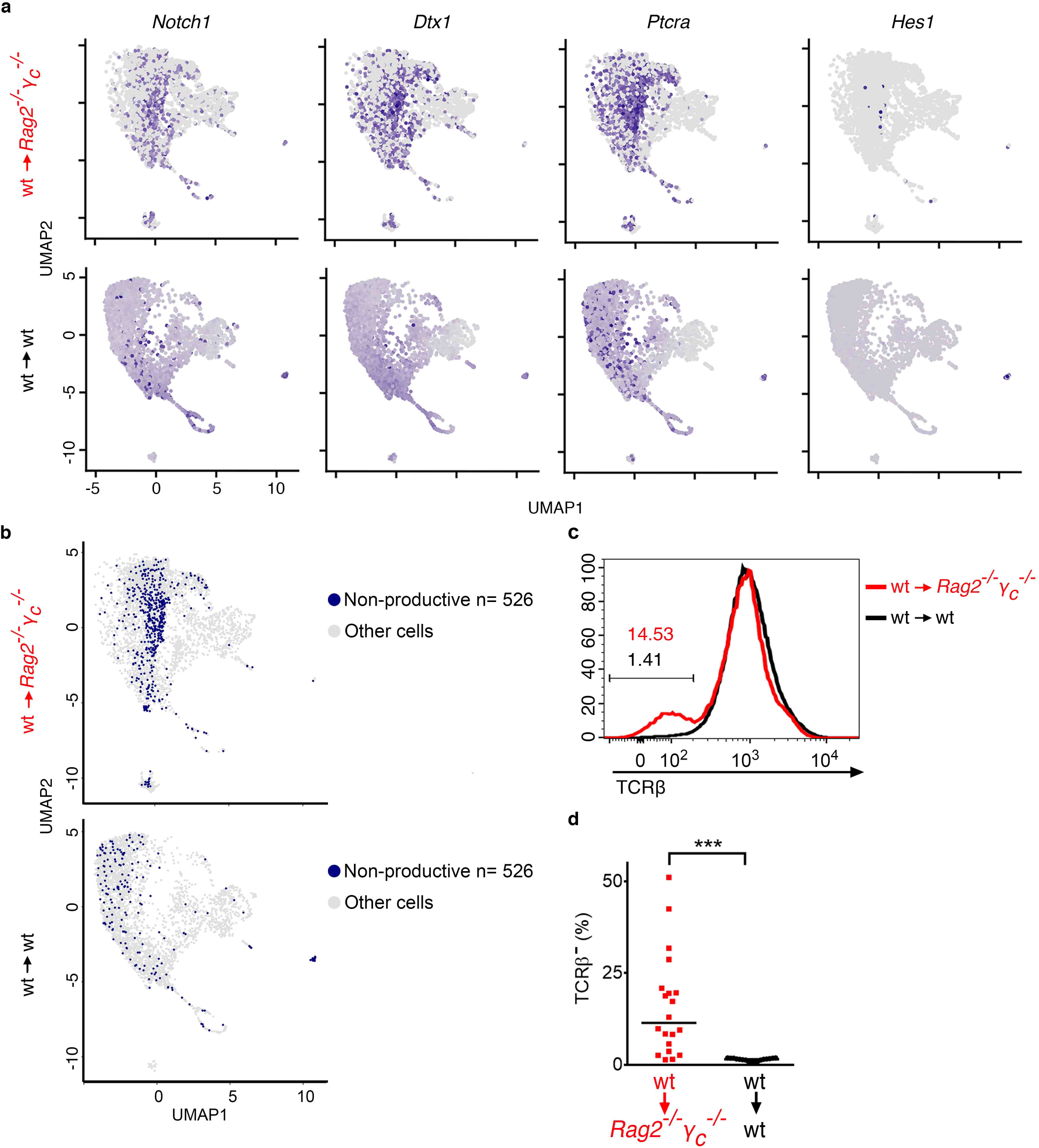
Emergence of aberrant thymocytes in thymus autonomy. **a)** UMAPs depict the expression of *Notch1* and selected Notch-target genes in thymus autonomy (top) or control thymocytes (bottom). **b)** UMAPs depict thymocytes with non-productive TCR in autonomy (top) or control thymocytes (bottom). **c)** Expression of TCRβ (intracellular staining) on CD4+CD8+ double positive thymocytes in autonomy (red) or control (black), 28 days after transplant. **d)** Quantification of the percentage of TCRβ-negative DP thymocytes in autonomy (red) or control (black) at day 28 after transplant. Each symbol represents one graft and the line marks the median. Data from 3 independent experiments. ***p<0.001

## DISCUSSION

Here we show that double negative 3 early (DN3e) thymocytes have the potential to self-renew and differentiate to maintain thymopoiesis autonomously if the thymus is deprived of competent bone marrow derived progenitors. While it remains unclear whether there is a physiological role for the self-renewal capacity of DN3e thymocytes, it is conceivable that they can maintain thymopoiesis following brief periods of progenitor deprivation. In this context it will be interesting to address whether models of deficiency in thymus homing, like the *Ccr7*^*-/-*^*Ccr9*^*-/-*^ mice^16, 17^, show a reduction in *TCRβ* diversity that could result from a short period of DN3e self-renewal. Similarly, conditions like infection, in which hematopoiesis is diverted from lympho- to myelopoiesis may also create a temporary shortage of lymphoid progenitors, and its impact could be attenuated by transient DN3e self-renewal. Of note, DN3 from autonomous thymi failed to grow on OP9-Dll4 cocultures that support T lymphocyte differentiation, suggesting that they require signals that differ from DN3 in the steady state, which proliferated and differentiated as expected (not shown).

Thymus autonomy, sustained by DN3e that self-renew and differentiate, is deleterious if prolonged because leukemia emerges as a final consequence^8, 10^. This highlights the importance of cell competition in the thymus, which occurs between DN2 to DN3e thymocytes and inhibits thymus autonomy^9^. Here we detected the emergence of aberrant cells as early as four weeks after induction of thymus autonomy. These are likely to be the cells giving rise to leukemia. The fact that the onset of T-ALL is at 15-week post-transplantation^8, 10^ suggests that leukemogenesis is a long process, and probably involves a series of events. Future studies will be important to detail the mechanisms that are active in the genesis of T-ALL following thymus autonomy.

An observation consistent with thymus autonomy prior to leukemia onset was reported following the successful correction of the human hematopoietic system by gene therapy in X-linked severe combined immunodeficiency (SCID-X1) in the early 2000’s^18^. SCID-X1 have loss-of-function mutations in the gene *γ*_*c*_, requiring a bone marrow transplant to survive. SCID-X1 has pioneered the first clinical trials of gene therapy for correction of a primary immunodeficiency^19^. Unfortunately, following gene therapy 5/20 of the patients developed T-ALL as a consequence^20, 21^. Malignant transformation was considered to result from genotoxicity due to vector integration close to oncogenes. While this may have been so, we have proposed that thymus autonomy could have contributed to this outcome^8^. In this context, two independent groups using bone marrow chimeras showed that the efficiency of bone marrow correction in this setting is an important factor to consider^22, 23^. If the reconstitution of the bone marrow is not efficient, then the efficiency of thymus seeding is likely to be compromised, leading to intermittent episodes of thymus seeding and function followed by periods refractory to thymus colonization. Following thymus seeding, thymopoiesis will be triggered. Provided that DN3e self-renew and maintain thymus autonomy, the main factor determining whether leukemia will eventually emerge will be the length of the period of thymus autonomy in between events of thymus colonization. Indeed, we consider that any condition enabling thymus autonomy for a prolonged time period may be permissive to T-ALL^24^.

## METHODS

### Mice

C57BL/6J (B6, CD45.2^*+*^) were bred and kept at the Instituto Gulbenkian de Ciência (IGC) in a colony that is refreshed frequently with mice purchased from Charles River. B6.SJL-*Ptprc*^a^ Pep3^b^/BoyJ (CD45.1^+^, here termed B6.SJL) mice, stock #002014, were purchased from The Jackson Laboratory and kept in a colony at IGC. Wild type donor thymi were harvested from newborn F1 donors (CD45.1+CD45.2+) resulting from the intercross between B6 and B6.SJL parents, unless stated otherwise. *Rag2*^*-/-*^*γ*_*c*_^*-/-*^ mice used here were imported from the University of Ulm, Germany, through embryo rederivation. *Rag2*^*-/-*^*γ*_*c*_^*-/-*^ mice in pure B6 background were obtained by crossing *γ*_*c*_^*-/-* 25^, JAX stock #003174 purchased from The Jackson Laboratory, and *Rag2*^*-/-* 26^, originally purchased from Taconic. *Rag2*^*-/-*^*γ*_*c*_^*-/-*^ are kept as two separate colonies but no differences were detected between using the strain imported, or the one in pure B6 background. Mice were kept in individually ventilated cages in specific pathogen-free (SPF) conditions and all experiments were conducted in compliance with Portuguese and European laws and approved by the Ethics Committee of the IGC – Fundação Calouste Gulbenkian and Direção Geral de Alimentação e Veterinária (DGAV).

### Thymus transplants

Thymic lobes from newborn donors were separated and grafted under the kidney capsule of recipient mice as described before, one lobe onto each extremity of the kidney ^6, 8^. Animals were anesthetized with Ketamine (100 mg/kg) and Xylazine (16 mg/kg) and B6 or *Rag2*^*-/-*^*γ*_*c*_^*-/-*^ mice aged 5 to 12 weeks were used as recipients (CD45.2^+^). Donors were F1:B6xB6.SJL (CD45.1^+^CD45.2^+^) in all experiments except for immunohistology shown in Fig.2, in which donors were B6.SJL (CD45.1^+^).

### Depletion of CD25-positive cells *in vivo*

Mice received a single injection of 250 μg of CD25-depleting antibody (PC61) or isotype IgG1 control (YCATE 55.9) i.p., 28 days after thymus transplant.

### Immunohistology

Thymus grafts were harvested and cryopreserved in OCT. 8 μm sections were obtained using a Leica Cryostat CM 3050 S, air dried, dehydrated in acetone and stored at -80°C until use. For staining, the slides were incubated with mouse IgG and DAPI, followed by primary antibodies overnight and secondary antibodies for 30 min. Antibodies used were anti-: rabbit Keratin 5 (Poly19055), Keratin 8-Alexa647 (1E8), CD11c-PE (N418), CD4-bio (GK1.5) and CD8-APC (53-6.7). Second step staining included Streptavidin-Cy3 and anti-rabbit a488. Slides were mounted using Fluromount G medium (Invitrogen) and images acquired using a Leica DMRA2 and a Nikon HCS microscope. Fiji was used for final image treatment.

### TEC isolation and staining

Thymic epithelial cells were isolated as previously^27^. In brief, minced grafts were digested on PBS containing 25 µg/ml of DNase I (Sigma-Aldrich), 100 µg/ml of Dispase I (Sigma-Aldrich) and 200 µg/ml of Collagenase/Dispase (Roche). Digestion was performed at 37 °C for 10 min for up to 4 rounds. Recovered supernatant were filtered through a 100 µm wide-pore mesh and ressuspended in PBS/10% FCS. Cells were blocked and then stained with antibodies recognizing CD45 biotin (30-F11), CD80 PerCP-Cy5.5 (16-10A1), EpCAM alexa647 (G8.8), and Ly51 PE (6C3), all from BioLegend.

### Flow cytometry and cell sorting

Organs were harvested and single-cell suspensions prepared in PBS/10% FBS. Cells were blocked with mouse IgG (Jackson laboratories) before staining. Antibodies and reagents purchased from BioLegend and recognizing the following antigens were: BrdU (3D4), CD3 (145-2C11), CD4 (GK1.1), CD8 (53-6.7; YTS169.8), CD11b (M1/70), CD11c (N418), CD19 (6D5), CD24 (M1/69), CD25 (PC61; 7D4), CD27 (LG.3A10), CD28 (37.51), CD44 (IM7), CD45.1 (A20), CD45.2 (104), CD71 (RI7217), CD98 (RL388), Gr-1 (RB6-8C5), KIT (2B8), Ki-67 (Ki67), NK1.1 (PK136), TCRβ (H57-597) and Ter119 (TER-119) conjugated to Alexa488, Alexa647, APC, APC-Cy7, APC-FIRE750, biotin, BV421, BV605, BV711, FITC, Pacific Blue, PE, PE-Cy7 and PercP-Cy5.5. Streptavidin conjugated to BV785 or APC-Cy7 was used to recognize biotinylated antibodies. The lineage cocktail consisted of CD3, CD4, CD8, CD11b, CD11c, CD19, GR-1, NK1.1 and Ter119. To assess apoptosis, after surface staining, thymocytes were incubated with Annexin V-PE (BioLegened), for 30 min at room temperature in Annexin binding buffer (0.15 M HEPES, 1.5 M NaCl and 0.25 M CaCl_2_). Samples were diluted in Annexin binding buffer and acquired immediately after staining. To assess glucose uptake, thymocytes were incubated in glucose-free RPMI with 100 μM 2-NBDG (Invitrogen) for 30 min at 37 °C with 5% CO2, before surface staining. Dead cells were excluded by sytox blue (Invitrogen) or zombie dyes (Pacific Orange or APC-Cy7, Biolegend). Intracellular staining was performed using True-Nuclear Transcription Factor Buffer Set (Biolegend). Samples were acquired using a BD LSRFortessa X-20 analyzer. Cell sorts were performed using a BD FACSAria II and sorted populations were identified as follows: DN3 (Lin^-^CD44^lo/-^CD25^+^), DN4 (Lin^-^CD44^-^CD25^-^CD45^+^), ISP (CD4^-^CD8^+^Lin^-^), DP (CD4^+^CD8^+^).

### EdU incorporation assays

Mice were injected intraperitoneally with 0.5 mg 5-ethynyl-2’-deoxyuridine (EdU, Invitrogen or Sigma-Aldrich). For pulse experiments, mice were injected twice (24 and 1 hours) or once (2 hours) before analysis, at specific timepoints after transplant, as indicated in the figures. For chase experiments, mice were injected twice, 6 hours apart, on day 28 after transplant. The first injection was considered day 0 and mice were analyzed at the timepoints indicated in the figure. For FACS analyses, cells were stained with extracellular antibodies followed by intracellular staining of EdU using click-iT Plus EdU Alexa Fluor 488 Flow Cytometry Assay Kit (Invitrogen), according to the manufacturer’s instructions.

### BrdU label retention assay

Transplanted mice were given BrdU in the drinking water (0.8mg/mL, Sigma-Aldrich), starting 14 days after transplant and continuously during 21 days. BrdU was replaced every two to three days. Thymus grafts were analyzed by flow cytometry at days 2 and 14 after BrdU withdrawal. For FACS analyses, extracellular antibody staining was performed followed by fixation in 4% PFA (15 minutes at room temperature) and permeabilization in saponin-based perm for 15 minutes at room temperature. Following a 1-hour incubation with DNAse (300ug/ml in PBS at 37°C, 5%CO2), cells were stained with an antibody cocktail containing anti-BrdU AlexaFluor488 and acquired in a BD LSRFortessa X-20 analyzer.

### Analysis of VDJ rearrangements of *TCRβ* in single cells

Single cells were sorted directly into 10 μl lysis buffer (1x Herculase II reaction buffer, 20 mg/ml Proteinase K (Thermofisher), 0.1 % Triton X100 (Sigma-Aldrich)) and digested (1 hour at 55 °C followed by 15’ at 95°C) as described elsewhere^28^. The first PCR was performed in a 20 μl reaction (1x Herculase II buffer, 50 nM each primer, 250 μM dNTPs and 0.4 μl Herculase II fusion DNA Polymerase). PCR reaction was 2 min at 98°C, followed by 35 cycles of 20 sec at 98°C, 20 sec at 66-58°C and 2 min at 72°C. The annealing temperature was decreased by 2 °C in each of the first 5 cycles. A final extension of 3 min was used. The second PCR was performed using 1 μl of the first PCR product in the same conditions. Primers used in both PCR rounds were the same as described^28^ (and supplementary table 5). Successful PCRs were confirmed by running 5 μl of the second PCR product in agarose gels. PCR products were purified and Sanger sequenced.

### Phasing of Sanger sequencing chromatograms

When 2 sequences from the same cell were detected, Sanger sequencing trace files were loaded to PolyPeakParser^29^ with a signal ratio cut-off of 0.3. This generated a heterozygous sequence (e.g. mixture of A and G is shown as R), which was loaded to a python script that searches for one PCR product in the mixture based on possible rearrangements of the *TCRβ* coding region. When the first sequence was resolved, it was given to PolyPeakParser as reference to generate the second sequence in the mixture. Furthermore, both sequences were aligned to GRCm38 TCRβ region for quality control.

### Bulk RNA-sequencing and data analysis

For each sample, 500 cells were sorted directly into Qiagen RLT lysis buffer. RNA was separated from genomic DNA using a modified oligo-dT bead-based method, adapted from ^30^. Libraries were prepared following the Pico Nextera protocol. Sequencing was performed using NextSeq500 high output kit (75 cycles), single-ended, 20 million reads per sample. Quality control was performed using FastQC. Samples were aligned using Hisat2 and counts obtained with Featurecount using the mouse genome mm10 GRCm38. Sequence alignments’ quality was evaluated using Qualimap. Differential gene expression analysis was conducted using the Deseq2 package in R (v3.6.1). Gene ontology enrichment analysis of the differentially expressed genes (with a corrected p-value bellow 0.05) was performed with Fisher exact text with Benjamini-Hochberg correction. Gene ontology terms with a corrected p-value bellow 0.01 were considered significant.

### 10x chromium Single cell gene expression and immune profiling

DN3-DN4, ISP and double positive thymocytes were sorted from one control thymus graft and from 4 pooled thymus grafts into PBS/10%BSA. Cell suspensions were loaded onto the Chromium Single cell Controller (10x Genomics) using Chromium Single Cell 5′ Library and the Gel Bead Kit following the manufacturer’s user guide (10x Genomics, Pleasanton, CA). The cDNA was amplified, using 10 PCR cycles, for downstream 5′ gene and enriched V(D)J library construction. The cDNA concentration was determined on an HS NGS Fragment Kit, Fragment Analyzer (Agilent). Libraries were constructed using the Chromium Single-Cell 5′ reagent kit, V(D)J Mouse T Cell Enrichment Kit, 5′ Library Construction Kit, and i7 Multiplex Kit (10x Genomics, CA) according to Chromium Single Cell 5′ Reagent Kit v2 user guide. Library quality was confirmed with an HS NGS Fragment Kit, Fragment Analyzer (Agilent) and Qubit Fluorometer (Thermo Fisher Scientific) with dsDNA HS assay kit according to the manufacturer’s recommendation. Sequencing of scRNA libraries: scRNAseq 5′ gene expression libraries and scRNAseq TCR V(D)J were loaded on an Illumina NextSeq 500 platform with a 500/550 High Output Kit v2 (300 cycles) to a read depth of ∼25,000 reads/cell and ∼7,500 reads/cell, respectively. All steps were performed at the Genomics Unit of the IGC.

### Single-cell RNA-seq raw data analysis

Illumina sequencer base call files (BCL) were processed using Cell Ranger mkfastq pipeline generating FASTQ files for both the 5′ expression library and the V(D)J libraries. Reads were mapped to the mouse genome build mm10. (available from 10X Genomics). The 5′ gene-expression libraries were then analyzed with the Cell Ranger count pipeline v3.0.2. The V(D)J data for each sample was processed using cellranger vdj command using the mouse reference dataset (GRCm38/Ensembl/10x) leading to the identification of the CDR3 sequences and the rearranged TCR genes. Analysis was performed using Loupe V(D)J Browser (v.3.0.0, 10x Genomics). In brief, a TCR diversity metric, containing clonotype frequency and barcode information, was obtained and Single cells were grouped in clonotypes when sharing the same TCRα and TCRβ sequences (V, D, J and CDR3 sequences).

### Single-cell RNA-seq analysis

The Seurat R package (v. 3.1.3.9002)^31^ was used to perform the quality-control, integration, dimensional reduction, clustering and exploration of control and autonomy scRNA-seq data sets. The matrix, gene (feature) and barcode tables provided by the upstream analysis of scRNA-seq data using the CellRanger pipeline were imported to Seurat with the Read10X function. The quality-control steps and filters applied were the same for the two data sets (control and autonomy): only genes expressed in at least 5 cells were kept; cells with more than 200 and less than 2500 genes were kept as well as cells with less than 5% of mitochondrial genes. Both data sets were independently normalized, using the LogNormalize method in Seurat with default options using the function NormalizeData. The 2,000 most variable genes were determined on each data set using the vst method with the FindVariableFeatures function. The total number of genes and the percentage of mitochondrial genes were regressed against each gene with the resulting residuals scaled and centered in order to minimize unwanted variation with the ScaleData function. The control and autonomy data sets were integrated by finding anchors between both data sets using 20 dimensions with the FindIntegrationAnchors and IntegrateData functions. The integrated data set comprising 2,000 genes and 7,488 cells (3,528 cells from control and 3,960 cells from autonomy data sets) was scaled before running the PCA (Principal Component Analysis) on the first 30 components with ScaleData and RunPCA functions, respectively. Two-dimensional UMAP (Uniform Manifold Approximation and Projection) was computed on the first 20 principal components based on the inspection of PCA result on the elbow plot. Then, clustering was performed using the first 20 principal components and a resolution of 0.4 using the FindNeighbors and FindClusters functions, respectively, yielding 11 cell clusters. Conserved gene markers between clusters were identified using the FindConservedMarkers function with a log fold-change threshold of 0.25 and only genes with an adjusted p-value <0.05, across both conditions (control and autonomy), were kept as conserved. Genes differentially expressed by cluster between the autonomy and control conditions were determined with the FindMarkers function applying a log fold-change threshold of 0.25 and an adjusted p-value <0.05. TCR (T-cell receptor) tables with annotation for control and autonomy scRNA-seq data sets were imported to R and the cell barcode was checked with those from gene expression tables that passed the filters applied through Seurat. This resulted in a match for 3,483 and 3,738 cells with TCR and gene expression information. Cells classified as “non-productive” were cells for which the column “raw_consensus_id” on the annotated TCR tables was equal to “None”. Cells with TCRα(s) and/or TCRβ(s) chain(s) were cells with a classification different from “None” in the column “raw_consensus_id” and classified as “TRA” and/or “TRB” in the column “chain”, respectively. The TCRα(s) or TCRβ(s) cdr3 sequences were retrieved checking the column “chain” for the alpha- or beta-TCR information and the cdr3 sequences retrieved from the column “cdr3”, respectively. Heatmaps were done with the ComplexHeatmap R package (v.2.0.0)^32^. All the downstream analyses mentioned previously were done on R version 3.6.2 or 3.6.3^33^.

### Statistical analysis

Datasets were tested for normality using D’Agostino & Pearson normality test. Mann-Whitney or t test with Welch’s correction were used as appropriate. Statistical analysis was conducted in Prism 7.

### Mathematical model of thymopoiesis

We developed a mathematical model for EdU incorporation during thymic development. We consider the DN3, DN4, ISP and DP compartments. For each compartment, we consider the (EdU) labeled and unlabeled (superscipt u) fractions. The model assumes that at the beginning of the experiment all compartments are EdU+ saturated, ie at their maximum level of labelling by EdU. Within each compartment, we consider cell division (*r*), cell death (*d*) and differentiation to the next developmental stage (*k*). Loss of EdU labelling is tied to cell division (α). The full model is described by the following equations:

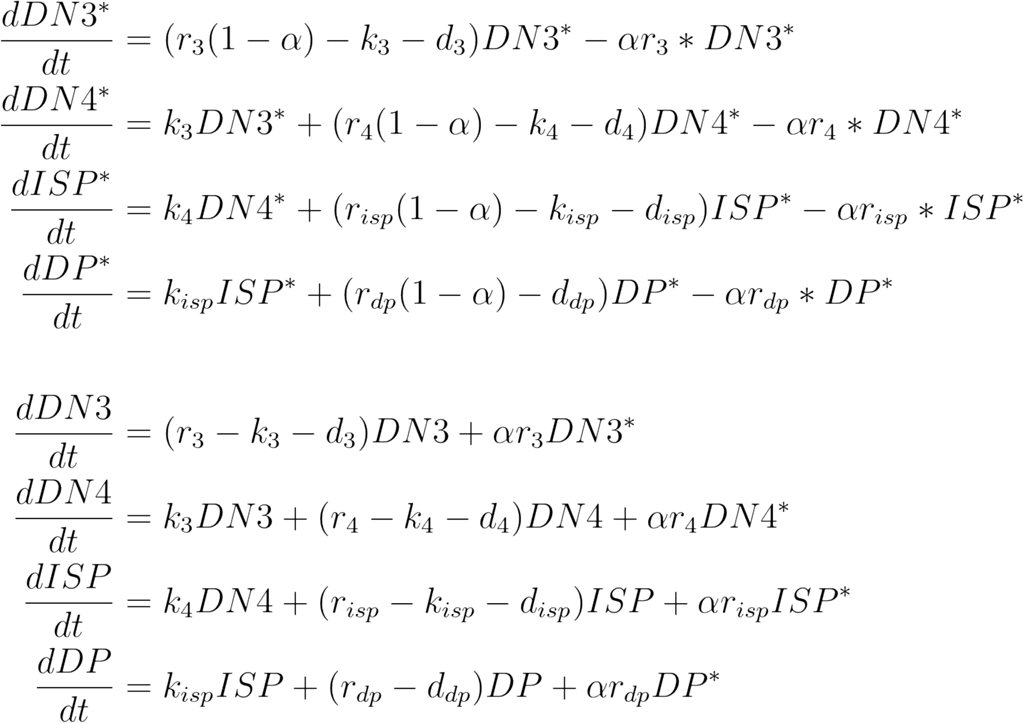

where * denotes EdU labelled cells.

The dynamics of the model was fit to the fraction of EdU+ cells within each compartment (eg DN3*/(DN3* + DN3)) by non-linear least squares regression (scipy 1.1). To account for the effect of the initial guesses for the values of the parameters, we sampled the parameter space uniformly to obtain a set of best-fit parameters. The quality of the fits was assessed by the Sum of Squared Residuals (SSR).

## ACKNOWLEDGEMENTS

This work was supported by the Instituto Gulbenkian de Ciência (IGC), Calouste Gulbenkian Foundation, and the Portuguese National Research Council (Fundação para a Ciência e Tecnologia [FCT] Grant PTDC/BIA-BID/30925/2017 to VCM). RAP and CVR are PhD students of the IGC Integrative Biology and Biomedicine (IBB) PhD Program and are supported by individual FCT PhD Fellowships refs. PD/BD/114341/2016 and PD/BD/139190/2018, respectively. This work had the support of the research infrastructures Congento LISBOA-01-0145-FEDER-022170 and PPBI-POCI-01-0145-FEDER-022122, both co-financed by FCT and Lisboa2020, under PORTUGAL2020 agreement (European Regional Development Fund). We acknowledge HJ Fehling and J Howard for critical reading of the manuscript. We acknowledge V Correia for technical support. We thank the team of Animal House Facility of IGC for the outstanding support to this work, and acknowledge the Microscopy, Antibody, Histopathology, Flow Cytometry, Genomics and Bioinformatics Units of IGC in supporting this work.

## AUTHOR CONTRIBUTIONS

RAP designed and performed experiments, analyzed data including the bioinformatic analysis of bulk RNAseq, and wrote the manuscript, AGGS performed the bioinformatic analysis of scRNAseq including the TCR diversity, CVR performed and analyzed the BrdU incorporation experiments, MA performed and analyzed the immunohistology, JL performed the phasing of sanger sequencing chromatograms when 2 overlapping sequences were detected, TP performed the mathematical modeling of the experiments involving EdU incorporation and chase, and VCM conceived the study, designed research, analyzed data including the preliminary bioinformatics scRNAseq and wrote the manuscript. All authors edited and contributed to the final version of the manuscript.

## REFERENCES

1. Rothenberg, E.V., Moore, J.E. & Yui, M.A. Launching the T-cell-lineage developmental programme. Nat Rev Immunol 8, 9–21 (2008).

2. Yui, M.A. & Rothenberg, E.V. Developmental gene networks: a triathlon on the course to T cell identity. Nat Rev Immunol 14, 529–545 (2014).

3. Klein, L., Kyewski, B., Allen, P.M. & Hogquist, K.A. Positive and negative selection of the T cell repertoire: what thymocytes see (and don’t see). Nat Rev Immunol 14, 377–391 (2014).

4. Boehm, T. Self-renewal of thymocytes in the absence of competitive precursor replenishment. J Exp Med 209, 1397–1400 (2012).

5. Frey, J.R., Ernst, B., Surh, C.D. & Sprent, J. Thymus-grafted SCID mice show transient thymopoiesis and limited depletion of V beta 11+ T cells. J Exp Med 175, 1067–1071 (1992).

6. Martins, V.C. et al. Thymus-autonomous T cell development in the absence of progenitor import. J Exp Med 209, 1409–1417 (2012).

7. Peaudecerf, L. et al. Thymocytes may persist and differentiate without any input from bone marrow progenitors. J Exp Med 209, 1401–1408 (2012).

8. Martins, V.C. et al. Cell competition is a tumour suppressor mechanism in the thymus. Nature 509, 465–470 (2014).

9. Ramos, C.V. et al. Cell competition, the kinetics of thymopoiesis and thymus cellularity are regulated by double negative 2 to 3 early thymocytes. bioRxiv doi.org/10.1101/2019.12.21.885673 (2020).

10. Ballesteros-Arias, L. et al. T Cell Acute Lymphoblastic Leukemia as a Consequence of Thymus Autonomy. J Immunol 202, 1137–1144 (2019).

11. Kelly, A.P. et al. Notch-induced T cell development requires phosphoinositide-dependent kinase 1. EMBO J 26, 3441–3450 (2007).

12. Williams, J.A. et al. Regulated costimulation in the thymus is critical for T cell development: dysregulated CD28 costimulation can bypass the pre-TCR checkpoint. J Immunol 175, 4199–4207 (2005).

13. Taghon, T., Yui, M.A., Pant, R., Diamond, R.A. & Rothenberg, E.V. Developmental and molecular characterization of emerging beta- and gammadelta-selected pre-T cells in the adult mouse thymus. Immunity 24, 53–64 (2006).

14. Benoist, C., Lanier, L., Merad, M., Mathis, D. & Immunological Genome, P. Consortium biology in immunology: the perspective from the Immunological Genome Project. Nat Rev Immunol 12, 734–740 (2012).

15. Mingueneau, M. et al. The transcriptional landscape of alphabeta T cell differentiation. Nat Immunol 14, 619–632 (2013).

16. Zlotoff, D.A. et al. CCR7 and CCR9 together recruit hematopoietic progenitors to the adult thymus. Blood (2010).

17. Liu, C. et al. Coordination between CCR7- and CCR9-mediated chemokine signals in prevascular fetal thymus colonization. Blood 108, 2531–2539 (2006).

18. Hacein-Bey-Abina, S. et al. Efficacy of gene therapy for X-linked severe combined immunodeficiency. N Engl J Med 363, 355–364 (2010).

19. Cavazzana-Calvo, M. et al. Gene therapy of human severe combined immunodeficiency (SCID)-X1 disease. Science 288, 669–672 (2000).

20. Howe, S.J. et al. Insertional mutagenesis combined with acquired somatic mutations causes leukemogenesis following gene therapy of SCID-X1 patients. J Clin Invest 118, 3143–3150 (2008).

21. Hacein-Bey-Abina, S. et al. Insertional oncogenesis in 4 patients after retrovirus-mediated gene therapy of SCID-X1. J Clin Invest 118, 3132–3142 (2008).

22. Schiroli, G. et al. Preclinical modeling highlights the therapeutic potential of hematopoietic stem cell gene editing for correction of SCID-X1. Sci Transl Med 9, eaan0820 (2017).

23. Ginn, S.L. et al. Limiting Thymic Precursor Supply Increases the Risk of Lymphoid Malignancy in Murine X-Linked Severe Combined Immunodeficiency. Mol Ther Nucleic Acids 6, 1–14 (2017).

24. Paiva, R.A., Ramos, C.V. & Martins, V.C. Thymus autonomy as a prelude to leukemia. FEBS J 285, 4565–4574 (2018).

25. Cao, X. et al. Defective lymphoid development in mice lacking expression of the common cytokine receptor gamma chain. Immunity 2, 223–238 (1995).

26. Shinkai, Y. et al. RAG-2-deficient mice lack mature lymphocytes owing to inability to initiate V(D)J rearrangement. Cell 68, 855–867 (1992).

27. Rode, I. et al. Foxn1 Protein Expression in the Developing, Aging, and Regenerating Thymus. J Immunol 195, 5678–5687 (2015).

28. Hosoya, T. et al. High-Throughput Single-Cell Sequencing of both TCR-beta Alleles. J Immunol 201, 3465–3470 (2018).

29. Hill, J.T. et al. Poly peak parser: Method and software for identification of unknown indels using sanger sequencing of polymerase chain reaction products. Dev Dyn 243, 1632–1636 (2014).

30. Macaulay, I.C. et al. Separation and parallel sequencing of the genomes and transcriptomes of single cells using G&T-seq. Nat Protoc 11, 2081–2103 (2016).

31. Stuart, T. et al. Comprehensive Integration of Single-Cell Data. Cell 177, 1888–1902 e1821 (2019).

32. Gu, Z., Eils, R. & Schlesner, M. Complex heatmaps reveal patterns and correlations in multidimensional genomic data. Bioinformatics 32, 2847–2849 (2016).

33. R_Core_Team. R: A language and environment for statistical computing.R Foundation for Statistical Computing, Vienna, Austria. https://www.R-project.org/ (2020).

