## SupplementaryFigures for "Self-renewal of double negative 3 (DN3) early thymocytes allows for thymus autonomy but compromises the β-selection checkpoint"

**Supplementary Figure 1**

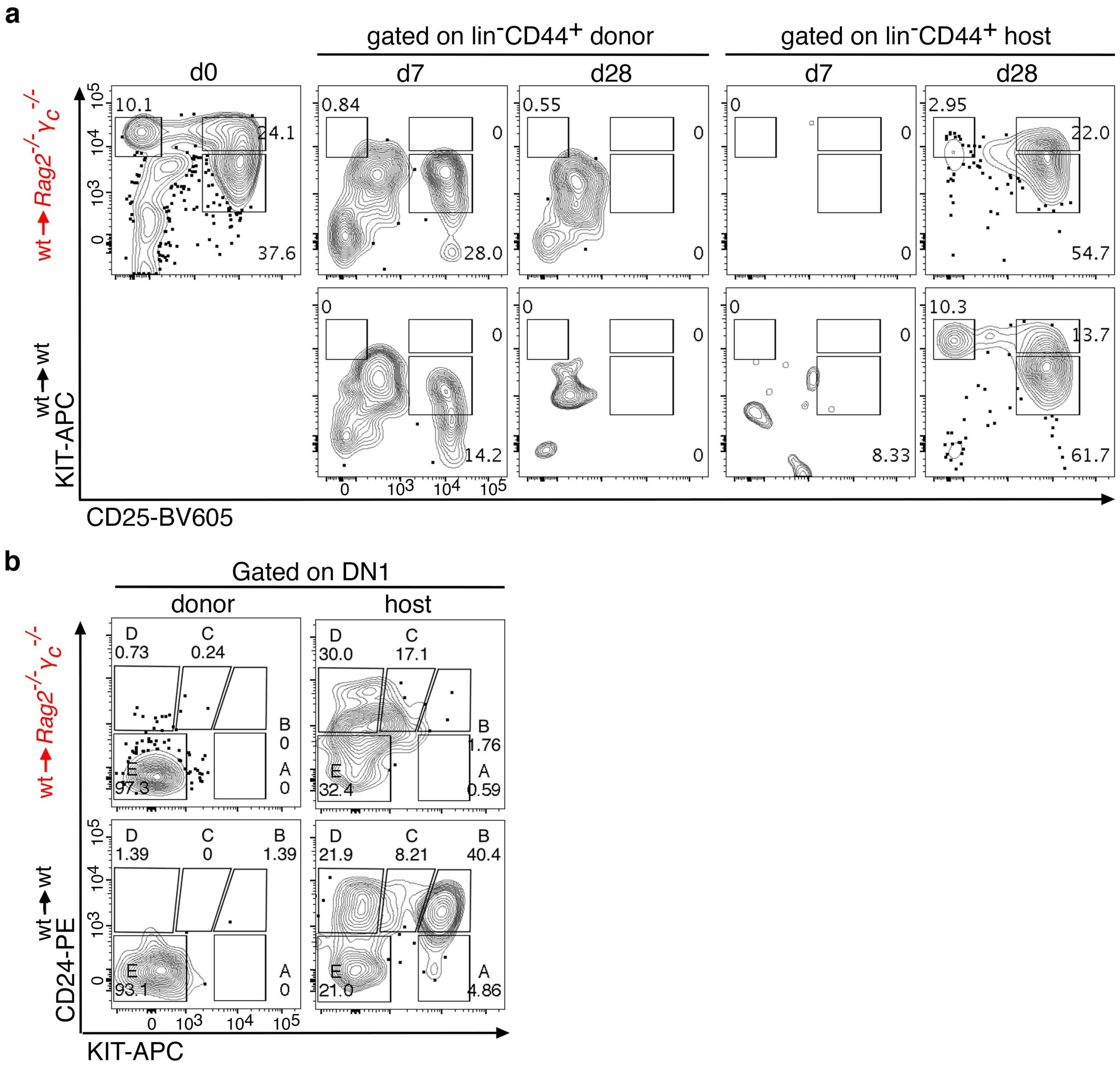

### Supplementary Figure 2

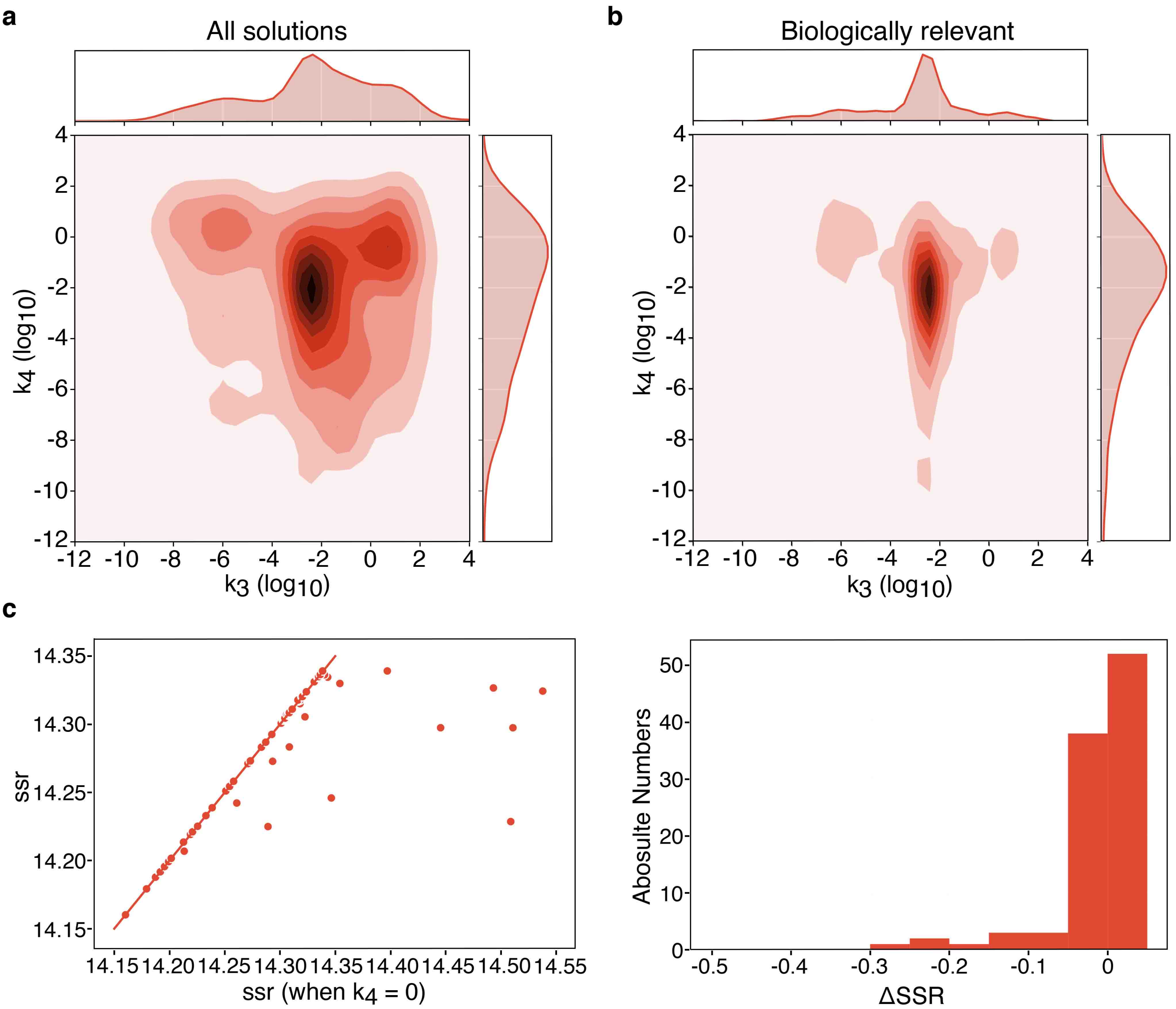

**Supplementary Figure 3**

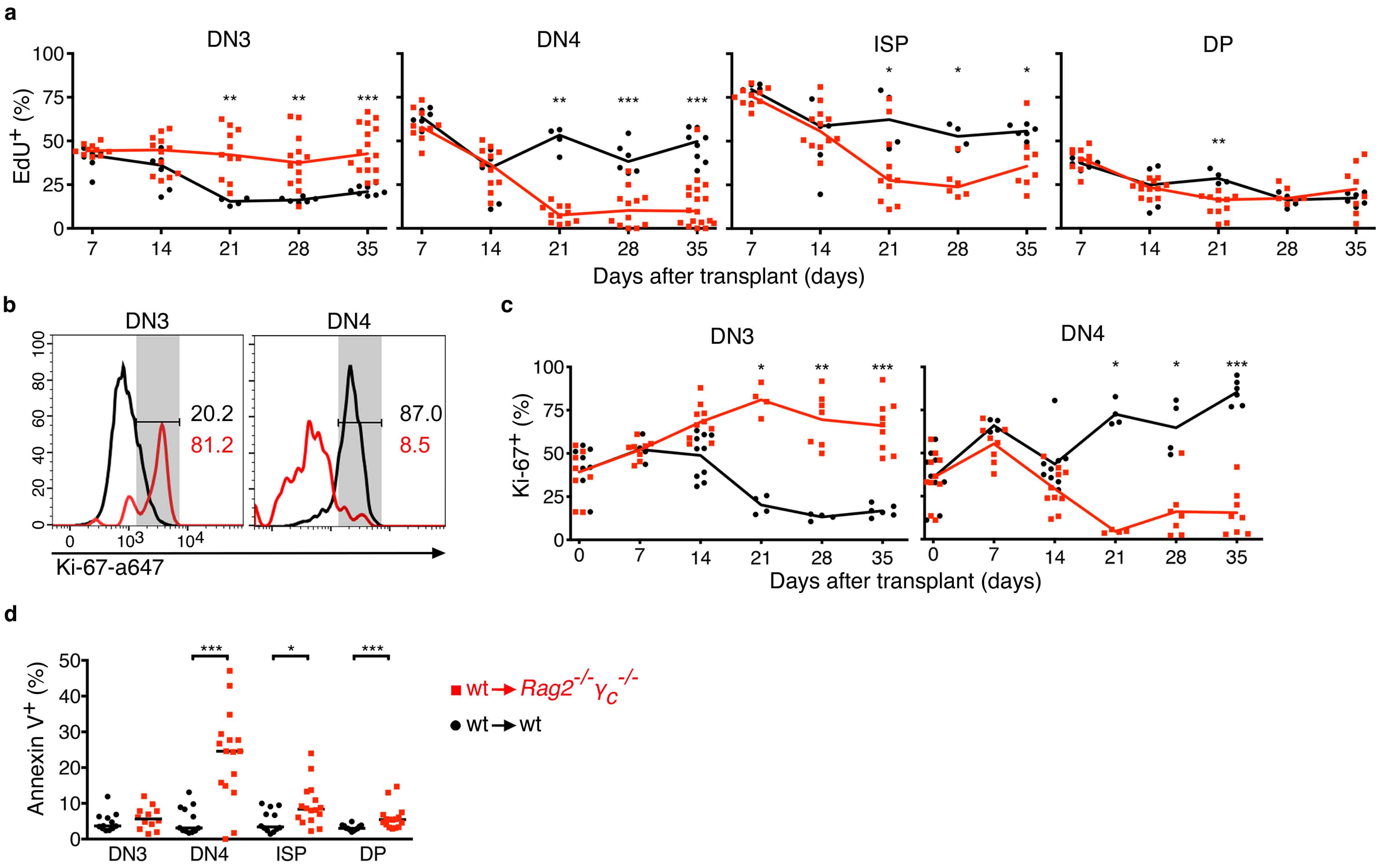

#### Supplementary Figure 4

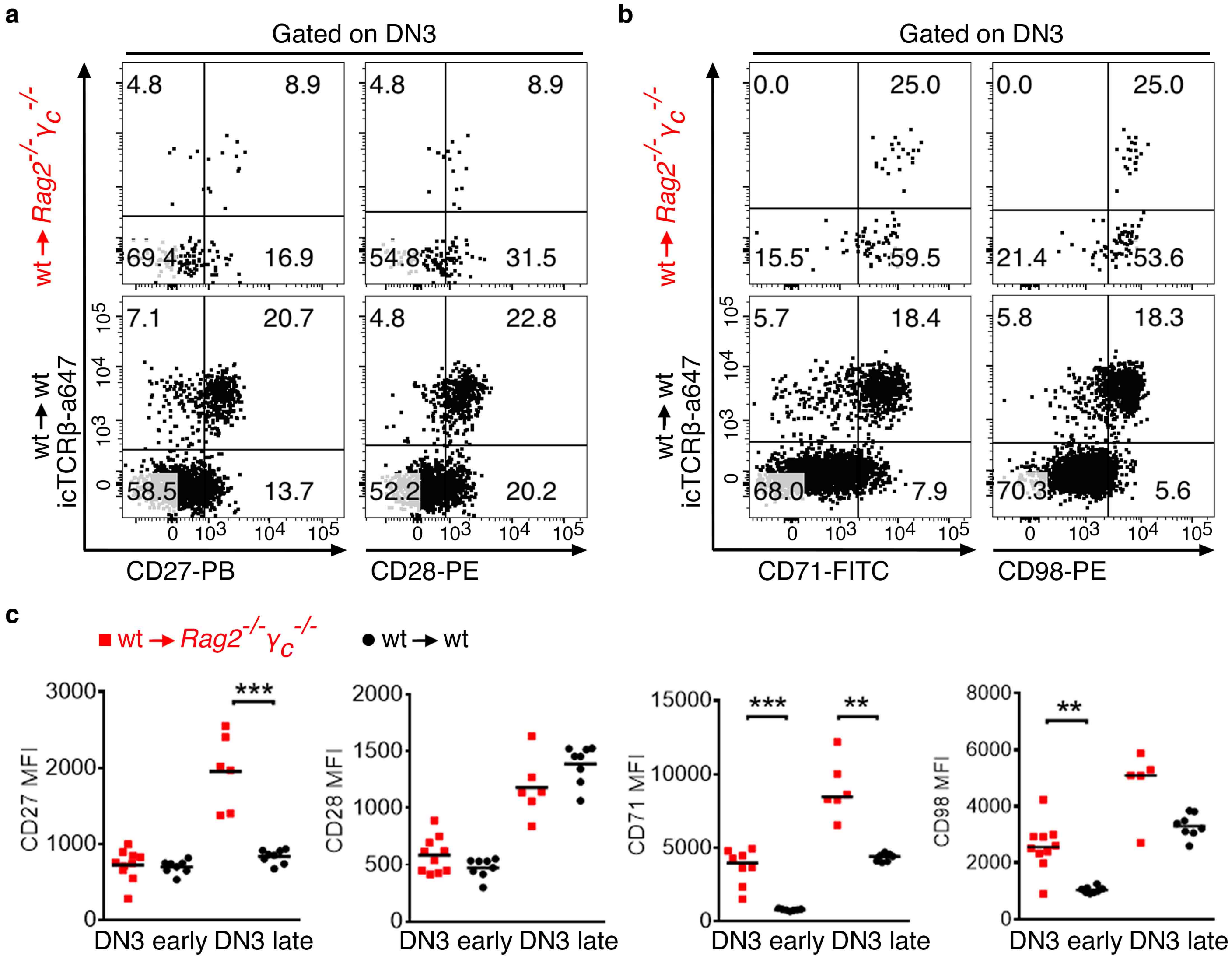

#### SUPPLEMENTARY FIGURE LEGENDS

**Supplementary Figure 1. Characterization of the most immature thymocytes in thymus transplants.** Thymus transplantation experiments were performed as depicted in Fig. 1a and analyzed by flow cytometry. **a)** Shown are representative examples analyzed at the indicated timepoints after transplant. Cells shown were gated on CD4-negative CD8-negative lineage-negative CD44-positive (DN1-2 compartment). One example is shown of a newborn thymus (day 0) and of grafts in autonomy (top) or control (bottom) 7 or 28 days after transplant. Cells were further pre-gated for donor or host origin, as indicated. **b)** Depicted is one example of a thymus graft in autonomy (top) or control (bottom) previously gated in thymocytes CD4-negative CD8-negative lineage-negative CD44-positive CD25-negative (DN1) 28 days after thymus transplant. Cells were further gated for donor (left) or host (right).

**Supplementary Figure 2. Modeling of differentiation in thymus autonomy. a-b)** Distribution of the differentiation rates from DN3 to DN4 (k3) and from DN4 to ISP (k4) obtained when no restrictions are applied to the model (a) or when we impose that the proliferation rates of DN3 and ISP thymocytes must be higher than that of DN4 and DP (b). **c)** Sum of squared residuals obtained from the original model (ssr) or when we test if no differentiation occurs between the DN4 and ISP stages (ssr when k4 = 0) and the distribution of the differences between the two conditions (right).

**Supplementary Figure 3. Autonomous thymopoiesis.** Wild type thymi were transplanted into *Rag2*$^{-/-}\gamma_c^{-/-}$ hosts and 6, 13, 20, 27, or 34 days later mice were injected i.p. with EdU once, and then again 24 hours later. Mice were analyzed 1 hour after the second injection. **a)** The indicated populations were analyzed for the percentage of EdU-positive cells at the indicated timepoints. Data was obtained from at least 2 independent experiments per timepoint. The data for DN3 at day 28 is the same shown in Fig. 4e. **b)** Thymocytes analyzed for Ki-67 expression 28 days after transplant, and shown are representative histograms in DN3 and DN4 thymocytes in thymus autonomy (gated on donor, red) or control (total, in black). **c)** Quantification of Ki-67 positive thymocytes at the indicated cells and timepoints. Each symbol is one graft and lines connect the median. Two independent experiments per timepoint, except for day 21 (n =1). **d)** Quantification of the percentage of Annexin V-positive thymocytes, in the indicated populations, at day 28 after transplant. Data obtained from 3 independent experiments. Each dot represents one graft and the lines mark the median. $^{*}p<0.05$, $^{**}p<0.01$, $^{***}p<0.001$

**Supplementary Figure 4. Immunophenotype of DN3 thymocytes in autonomy.** Thymus transplantation experiments were performed as depicted in Fig. 1a and analyzed by flow cytometry. **a-b)** One example of thymocytes 28 days after transplant gated on donor (thymus autonomy, top) or total (control, bottom) DN3 for **a)** icTCRβ versus CD27 or CD28, and **b)** and icTCRβ versus CD71 or CD98. **c)** Quantification of the mean fluorescence intensity (MFI) of the indicated markers in DN3e and DN3l in autonomy (red) or control (black). Each dot represents one graft and the lines mark the median. Data from 2 independent experiments. $^{**}p<0.01$, $^{***}p<0.001$
